# m6A-dependent circular RNA formation mediates tau-induced neurotoxicity

**DOI:** 10.1101/2024.01.25.577211

**Authors:** Farzaneh Atrian, Paulino Ramirez, Jasmine De Mange, Marissa Marquez, Elias M. Gonzalez, Miguel Minaya, Celeste M. Karch, Bess Frost

**Affiliations:** Sam & Ann Barshop Institute for Longevity and Aging Studies; Glenn Biggs Institute for Alzheimer’s and Neurodegenerative Diseases; Department of Cell Systems and Anatomy; University of Texas Health San Antonio, San Antonio, TX; Department of Psychiatry, Washington University, St Louis, MO

**Keywords:** Alzheimer’s disease, tauopathy, circRNA, Mbl, RNA methylation, nucleus

## Abstract

Circular RNAs (circRNAs), covalently closed RNA molecules that form due to back-splicing of RNA transcripts, have recently been implicated in Alzheimer’s disease and related tauopathies. circRNAs are regulated by N^6^-methyladenosine (m^6^A) RNA methylation, can serve as “sponges” for proteins and RNAs, and can be translated into protein via a cap-independent mechanism. Mechanisms underlying circRNA dysregulation in tauopathies and causal relationships between circRNA and neurodegeneration are currently unknown. In the current study, we aimed to determine whether pathogenic forms of tau drive circRNA dysregulation and whether such dysregulation causally mediates neurodegeneration. We identify circRNAs that are differentially expressed in the brain of a *Drosophila* model of tauopathy and in induced pluripotent stem cell (iPSC)-derived neurons carrying a tau mutation associated with autosomal dominant tauopathy. We leverage *Drosophila* to discover that depletion of circular forms of *muscleblind* (*circMbl)*, a circRNA that is particularly abundant in brains of tau transgenic *Drosophila*, significantly suppresses tau neurotoxicity, suggesting that tau-induced *circMbl* elevation is neurotoxic. We detect a general elevation of m^6^A RNA methylation and circRNA methylation in tau transgenic *Drosophila* and find that tau-induced m^6^A methylation is a mechanistic driver of *circMbl* formation. Interestingly, we find that circRNA and m^6^A RNA accumulate within nuclear envelope invaginations of tau transgenic *Drosophila* and in iPSC-derived cerebral organoid models of tauopathy. Taken together, our studies add critical new insight into the mechanisms underlying circRNA dysregulation in tauopathy and identify m^6^A-modified circRNA as a causal factor contributing to neurodegeneration. These findings add to a growing literature implicating pathogenic forms of tau as drivers of altered RNA metabolism.

## INTRODUCTION

Alzheimer’s disease is a progressive neurodegenerative disorder and the most common cause of dementia (Knopman et al., 2021). The neuropathological hallmarks of Alzheimer’s disease, amyloid beta plaques and tau tangles, form decades prior to cognitive decline (Braak et al., 2011). In addition to Alzheimer’s disease, tau aggregates are a defining feature of diverse neurodegenerative disorders collectively known as “tauopathies,” some of which arise due to mutations in the *MAPT* gene (Hutton et al., 1998; Poorkaj et al., 1998; Spillantini et al., 1998), which encodes tau protein. Various models of tauopathy that rely on transgenic expression of frontotemporal dementia-associated *MAPT* mutants have provided novel insights into the basic biology underlying human Alzheimer’s disease and related tauopathies (Khurana, 2008; Jankowsky & Zheng, 2017), despite the fact that most primary and secondary tauopathies are sporadic and thus involve the deposition of wild-type forms of tau. Studies in *Drosophila* models of tauopathy indicate that pathogenic forms of wild-type tau and tau harboring various frontotemporal dementia-associated mutations drive neurotoxicity through a common pathway involving the actin cytoskeleton and consequent changes in nuclear and genomic architecture (Frost et al., 2014; Frost et al., 2016; Bardai et al., 2018).

We and others have previously reported that wild-type and mutant forms of tau drive neurodegeneration by negatively affecting the overall three-dimensional structure of the nucleus. Neurons from brains of tau transgenic *Drosophila,* human brain affected by Alzheimer’s disease (Frost et al., 2016) frontotemporal dementia due to *MAPT* mutation, iPSC-derived neurons carrying a *MAPT* mutation (Paonessa et al., 2019), cultured HEK293 or neuroblastoma cells with induced tau expression (Montalbano et al., 2020; Sohn et al., 2023), and primary cultured cortical neurons with optogenetically-induced tau multimerization (L. Jiang et al., 2021) harbor nuclear invaginations and/or blebs. Studies in tau transgenic *Drosophila* suggest that such nuclear pleomorphisms are caused by destabilization of the lamin nucleoskeleton and causally mediate neuronal death (Frost et al., 2016).

We have previously discovered that polyadenylated (polyA) RNA accumulates within tau-induced nuclear invaginations and blebs in the *Drosophila* brain, and that such accumulation can be reduced by genetic or pharmacologic reduction of RNA export, which also suppresses tau-induced neurotoxicity (Cornelison et al., 2019). Along with studies showing that pathogenic forms of tau trigger and/or are associated with abnormal RNA splicing and intron retention (Apicco et al., 2019; Hsieh et al., 2019; Koren et al., 2020), we recently found that clearance of aberrant RNA transcripts via nonsense-mediated mRNA decay (NMD) is blunted in tauopathy, and that proteins translated from RNA carrying NMD-triggering features also accumulate in tau-induced nuclear blebs (Zuniga et al., 2022). There is thus a growing literature documenting alterations in RNA metabolism in tauopathy, and association of such RNA species with tau-induced nuclear pleomorphisms.

We became interested in the potential role of tau as a mediator of circRNA biogenesis based on previous findings that circRNAs are differentially expressed in human Alzheimer’s disease brain tissue (Dube et al., 2019). CircRNA biogenesis occurs through a back-splicing event that is independent of the parental linear RNA (Salzman et al., 2013; Knupp & Miura, 2018). With stable structure and long half-life in cerebrospinal fluid, circRNAs are hallmarks of aging that have been proposed as potential biomarkers for neurodegenerative disorders (Memczak et al., 2015; Knupp & Miura, 2018; Dube et al., 2019). Mechanistically, circRNA can serve as a sponge for complementary RNAs and RNA binding proteins (Hansen et al., 2013; Memczak et al., 2013; Gokool et al., 2020). In addition, recent work suggests that some circRNAs can be actively translated into protein via a cap-independent translation mechanism (Pamudurti et al., 2017; Yang et al., 2017). CircRNA formation, subsequent export from the nucleus, and, in some cases, translation into protein, are regulated in part by m^6^A modification (Yang et al., 2017; Di Timoteo et al., 2020), the most abundant RNA modification in eukaryotes. While m^6^A modified RNA is elevated in human Alzheimer’s disease and in tau transgenic mice (Han et al., 2020; L. Jiang et al., 2021), circRNA is dysregulated in Alzheimer’s disease, and m^6^A is known to regulate circRNA formation, the link between tau, m6A, circRNA and neurotoxicity have yet to be explored.

In the current study, we find that *circMbl* is significantly elevated as a consequence of transgenic panneuronal expression of a frontotemporal dementia-associated human *MAPT* mutation (tau^R406W^) in the adult *Drosophila* brain. Genetic reduction of *circMbl* suppresses tau-induced neuronal death, suggesting that *circMbl* mediates tau neurotoxicity. Mechanistically, we find that tau-induced elevation of RNA methylation regulates *circMbl* abundance in brains of tau transgenic *Drosophila* and observe that circRNA and m^6^A closely associate with nuclear blebs that form in tau transgenic *Drosophila* and in iPSC-derived cerebral organoids from a patient harboring a *MAPT* mutation (tau^IVS10+16^). Taken together, our studies reveal a previously unknown link between tau, RNA methylation, circRNA biogenesis, and neurodegeneration, and point toward the potential involvement of nuclear blebs as a nuclear export system for accumulating circRNA.

## RESULTS

### CircMbl is elevated in the adult brain of a *Drosophila* model of tauopathy

Formation of aberrantly phosphorylated, misfolded tau species lead to neuronal death and cognitive decline in various laboratory models of tauopathy, including *Drosophila*. We used a well-described *Drosophila* model of tauopathy (Wittmann et al., 2001) to determine if the differential expression of circRNAs observed in human Alzheimer’s disease cases was conserved in this model. Pan-neuronal expression of human tau carrying the frontotemporal dementia-associated *R406W* (Hutton et al., 1998) mutation in the adult *Drosophila* brain causes age-associated neuronal death, shortened lifespan, and decreased locomotor activity (Wittmann et al., 2001; Frost et al., 2014). This model recapitulates many cellular features of human Alzheimer’s disease and related tauopathies, including but not limited to aberrant tau phosphorylation, oxidative stress (Dias-Santagata et al., 2007), DNA damage (Khurana et al., 2012), and nuclear envelope invaginations and blebs (Frost et al., 2016).

We utilized the circRNA analysis tool DCC (Cheng et al., 2016) to identify differentially expressed circRNAs (**Supplemental Dataset 1**) in a publicly available RNA-seq dataset from heads of control and tau^R406W^ transgenic *Drosophila* at day 10 of adulthood (Mahoney et al., 2020), an age at which neuronal death and locomotor deficits are detectable in this model but prior to significant decline in survival (Frost et al., 2014). CircRNA analysis algorithms recognize the back-splice junction that forms between the 5’ and 3’ ends of exons when RNAs circularize (**Fig. 1A**). We identify circular forms of *muscleblind* (*mbl*) and *beaten path Vc* (*beat-Vc*) as significantly elevated in heads of tau^R406W^ transgenic *Drosophila* (**Fig. 1B**). As muscleblind is known for its role in splicing, transcript localization, and miRNA/circRNA biogenesis (Houseley et al., 2006), and has previously been implicated in various neurological disorders (Kanadia et al., 2003; de Haro et al., 2006; Rudnicki et al., 2007; Li et al., 2008; Daughters et al., 2009; Sellier et al., 2010; Casci et al., 2019), we focused on *circMbl* (**Fig. 1C**) for subsequent mechanistic analyses.

**Figure 1.**
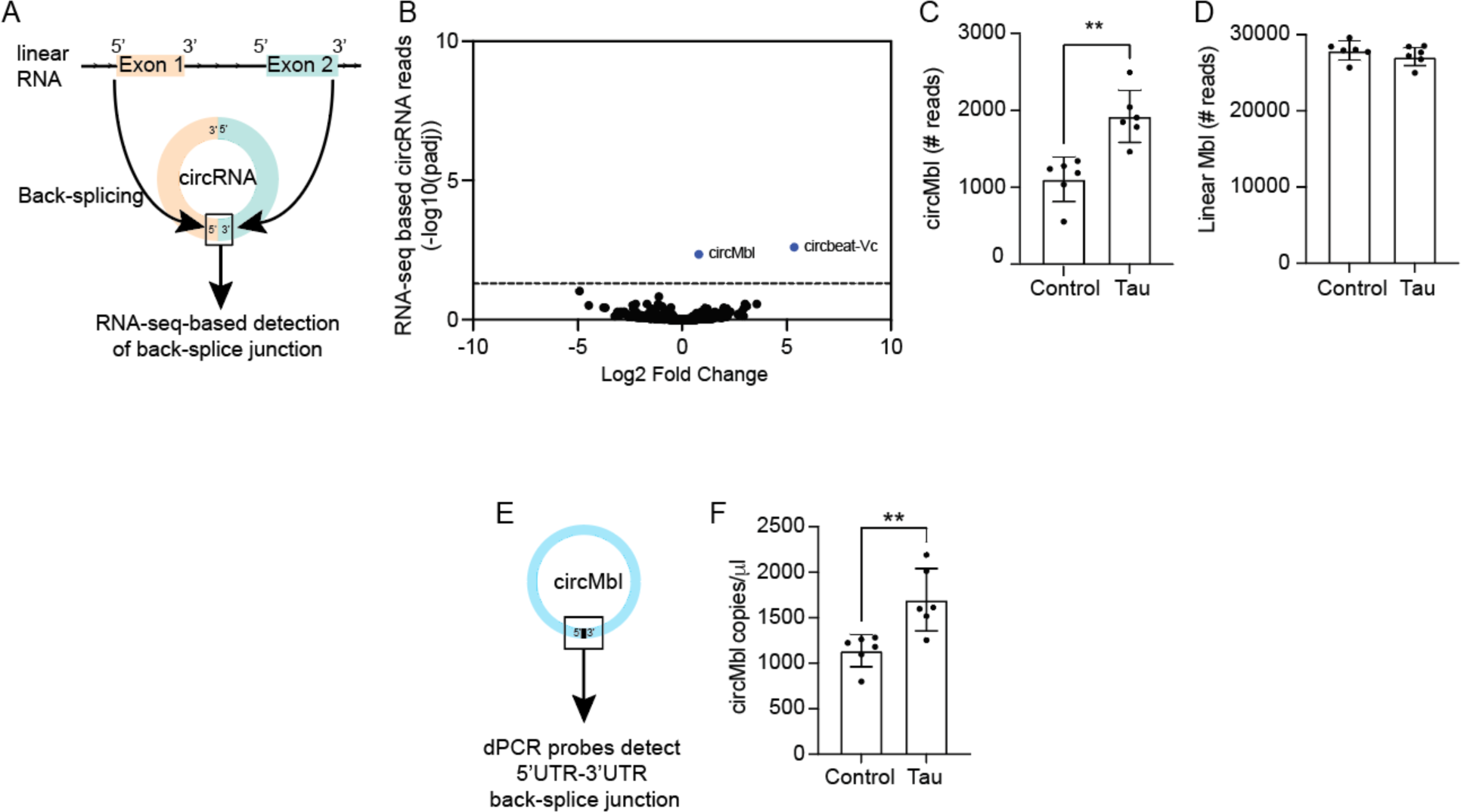
Increase in circRNA abundance in heads of adult tau^R406W^ transgenic *Drosophila*. **A**. Cartoon depicting formation of circRNA from a linear transcript. Back-splice junction indicates a circRNA. **B**. Differentially expressed circRNAs in tau^R406W^ transgenic *Drosophila* heads compared to control based on DCC RNA-seq analysis. Dashed line indicates an adjusted p value of 0.05. **C**. *CircMbl* in tau^R406W^ transgenic *Drosophila* heads is significantly increased compared to control based on DCC analysis. **D**. RNA-seq based reads for linear *mbl*. **E**. Cartoon depicting the 5’-3’ back-splice junction of *circMbl* used to design dPCR probes. **F**. *CircMbl* copies are significantly elevated in tau^R406W^ transgenic *Drosophila* compared to control based on dPCR. *n* = 6 biological replicates per genotype. \*\**p* < 0.01, unpaired t-test. Error bars = SEM. All analyses were performed at day 10 of adulthood.

RNA-seq reveals that levels of linear *mbl* transcript are unchanged in tau^R406W^ transgenic *Drosophila* versus controls (**Fig. 1D**), consistent with previous reports that circRNA levels are independent of their linear RNA counterparts (Ashwal-Fluss et al., 2014). We next designed digital PCR (dPCR) probes to specifically detect the 5’-3’ back-splice junction of *circMbl* (**Fig. 1E**). dPCR of RNA extracted from control and tau^R406W^ transgenic heads confirms that this circRNA species is significantly elevated in tau^R406W^ transgenic *Drosophila* (**Fig. 1F**). Taken together, our data suggests that pathogenic forms of tau drive an increase in circular, but not linear, *mbl* transcripts.

### Reduction of *circMbl* suppresses tau-induced neurotoxicity in *Drosophila*

Having found that *circMbl* is significantly elevated in brains of tau transgenic *Drosophila,* we next asked if tau-induced elevation of *circMbl* causally drives neurotoxicity. We reduced *circMbl* and total *mbl* in neurons of tau transgenic *Drosophila* using two approaches: 1) Targeted depletion of *circMbl* via pan-neuronal expression of a small hairpin RNA that is complementary to the exon 2 back-splice junction in *circMbl* (*circMbl^KD^*) (Pamudurti et al., 2022) and 2) General pan-neuronal RNAi-mediated knockdown of *mbl* (*mbl^RNAi^*), which targets linear *mbl* and *circMbl*. We find that both *circMbl^KD^* and *mbl^RNAi^* significantly deplete *circMbl* in brains of tau transgenic *Drosophila* based on dPCR (**Fig. 2A**). To further confirm that *circMbl* is reduced in the context of *circMbl^KD^* and *mbl^RNAi^*, we designed fluorescence *in situ* hybridization (FISH) probes to detect the 5’-3’ back-splice junction of *circMbl*. Indeed, we observe significantly less *circMbl* in brains of tau transgenic flies with *circMbl^KD^*or *mbl^RNAi^*, consistent with dPCR (**Fig. 2B, C**).

**Figure 2.**
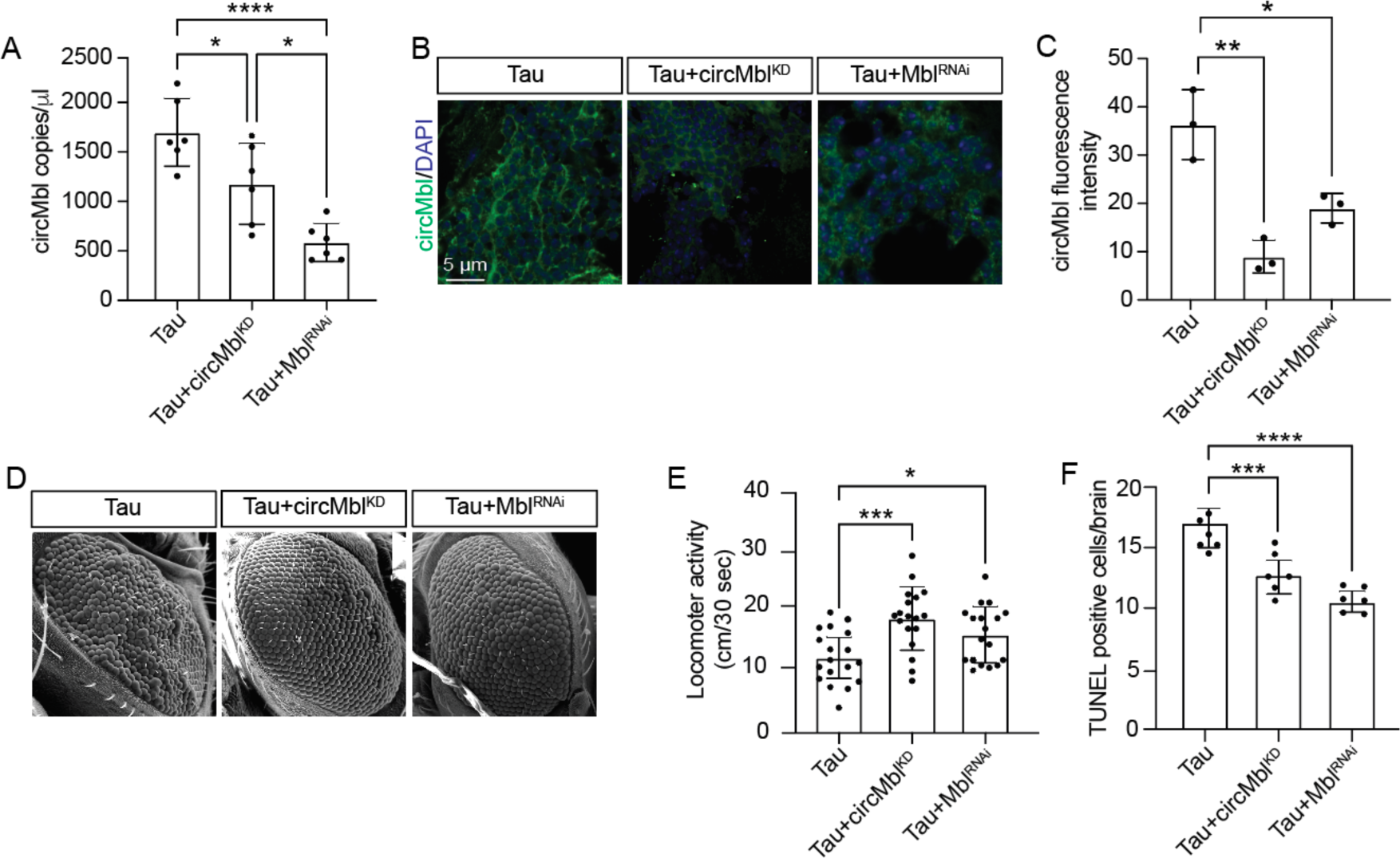
Reduction of *circMbl* suppresses tau-induced neurotoxicity in the *Drosophila* brain. **A**. Panneuronal expression of *circMbl^KD^ or mbl^RNAi^* significantly reduce *circMbl* in heads of tau transgenic *Drosophila* based on dPCR. **B**. FISH-based detection of *circMbl* in tau, tau+*circMbl*^KD^ and tau+*Mbl^RNAi^ Drosophila* brains. **C**. Quantification of (B). **D**. *circMbl^KD^* and *mbl^RNAi^* suppress the tau-induced rough eye phenotype. **E**. *circMbl^KD^* and *mbl^RNAi^*alleviate tau-induced locomotor deficits. **F**. *CircMbl^KD^* and *mbl^RNAi^* suppress tau-induced neurodegeneration based on TUNEL. *n* = 6 or 18 biological replicates per genotype. \**p* < 0.05, \*\**p* < 0.01, \*\*\**p* < 0.001, \*\*\*\**p* < 0.0001, ANOVA with a post-hoc Tukey test was used for multiple comparisons. Error bars = SEM. All analyses were performed at day 10 of adulthood.

Having established that *circMbl^KD^* and *mbl^RNAi^* significantly deplete overall levels of *circMbl* in brains of tau transgenic *Drosophila,* we next determined if tau-induced elevation of *circMbl* causally mediates neurodegeneration using three different commonly-used assays of tau-induced neurotoxicity in *Drosophila*. We find that *circMbl^KD^* and *mbl^RNAi^* suppress the tau-induced rough eye phenotype (**Fig. 2D**), and significantly improve tau-induced deficits in locomotor activity (**Fig. 2F**). To quantify neurodegeneration in the brain directly, we utilized the Terminal deoxynucleotidyl Transferase (TdT) dUTP Nick-End Labeling (TUNEL) assay, which labels DNA damage associated with apoptotic cell death (Kyrylkova et al., 2012). We find that depletion of *circMbl* or *mbl* significantly reduces apoptotic cell death in brains of tau transgenic *Drosophila* (**Fig. 2G**), and slightly but significantly reduces levels of tau protein (**Supplemental Fig. 1A, B**).

### RNA methylation mediates tau-induced RNA circularization

Having identified circRNAs that are elevated in brains of tau transgenic *Drosophila,* and that tau-induced elevation of *circMbl* causally mediates neurotoxicity, we next investigated the mechanism underlying tau-induced *circMbl* elevation in tau transgenic *Drosophila.* It is now well understood that RNA is subject to epigenetic modification, and that such modifications regulate RNA alternative splicing, circularization, stability, nuclear export, and translation (X. Jiang et al., 2021). As m^6^A is the most common epigenetic modification of RNA (Dubin & Taylor, 1975; Perry et al., 1975) and is reported to regulate RNA circularization (Zhou et al., 2017), we investigated RNA methylation as a candidate mechanism regulating tau-induced elevation of circRNA. We first analyzed overall levels of m^6^A in total head lysates from tau transgenic *Drosophila*. Based on dot blot, we find a significant increase in total m^6^A levels in the *Drosophila* model of tauopathy compared to control (**Fig. 3A, B**). To visualize m^6^A in the brain directly, we performed m^6^A immunofluorescence in brains of control and tau transgenic *Drosophila.* These analyses further indicate that m^6^A is significantly elevated in the context of tauopathy (**Fig. 3C, D**).

**Figure 3.**
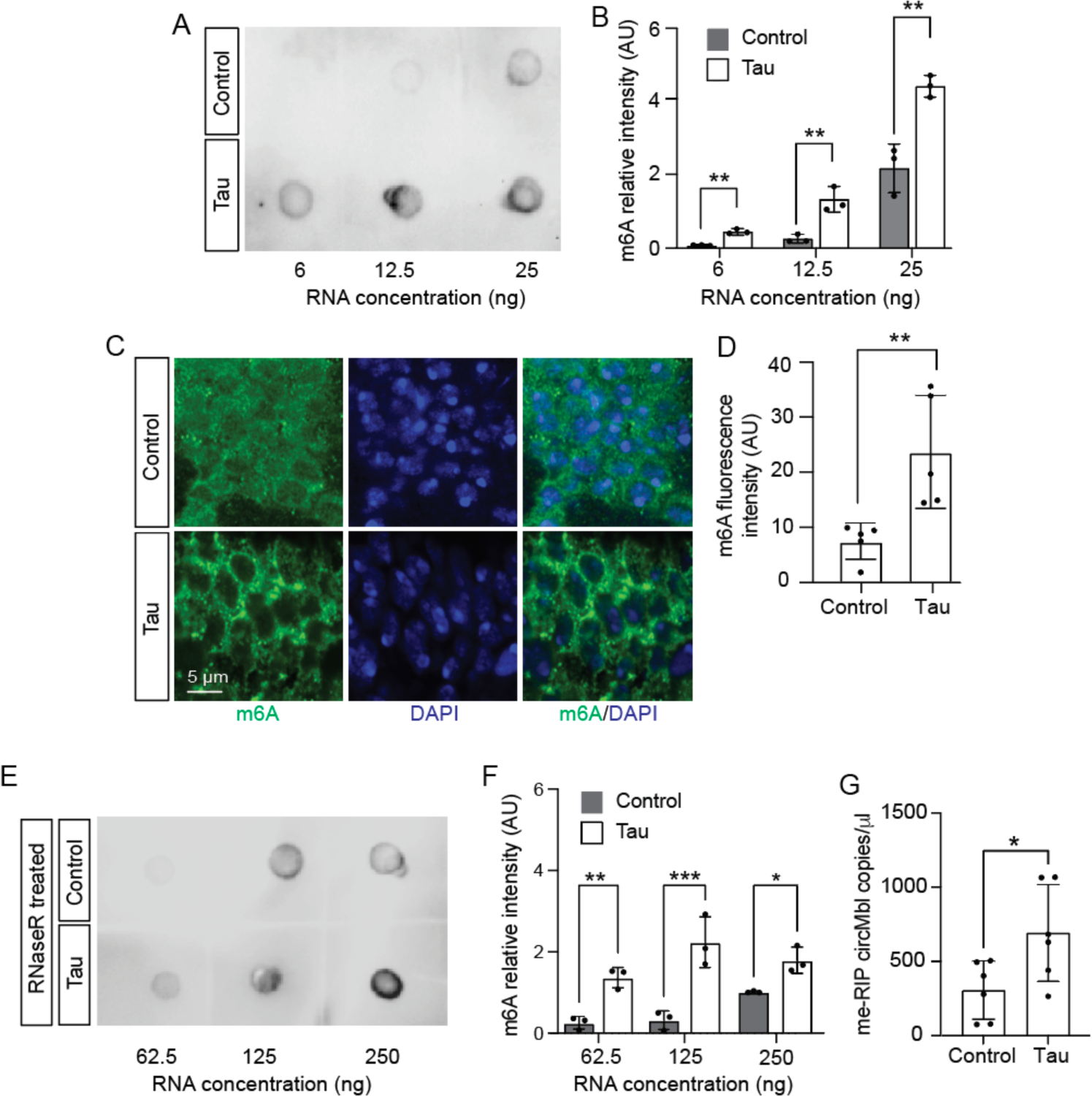
m^6^A mediates tau-induced circularization of *mbl* in brains of tau transgenic *Drosophila*. **A**. Levels of m^6^A in total RNA from control and tau transgenic *Drosophila* based on dot blot. RNA was loaded to the blot at increasing concentrations for each genotype. **B**. Quantification of (A). **C**. Visualization of m^6^A in brains of control and tau transgenic *Drosophila* based on immunofluorescence. **D**. Quantification of (C). **E**. Levels of m^6^A in RNase R-treated RNA samples from control and tau transgenic *Drosophila* based on dot blot. RNA was loaded to the blot at increasing concentrations for each genotype. **F**. Quantification of (E). **G**. *CircMbl* copies in m^6^A enriched RNA (me-RIP) samples from control and tau^R406W^. n = 6 biological replicates per genotype. *p<0.05, **p,0.01, ***p<0.001, unpaired t-test. Error bars = SEM. All flies were analyzed at day 10 of adulthood.

We next determined the degree to which circRNAs are methylated in head lysates from control and tau transgenic *Drosophila.* We first enriched for circRNA by digesting linear RNA with Ribonuclease R, then subjected remaining circRNA to m^6^A analysis via dot blot. We find that circRNAs are m^6^A modified to a greater extent in tau transgenic *Drosophila* compared to control (**Fig. 3E, F**). To investigate the m^6^A status of *circMbl* specifically, we performed EpiQuik CUT&RUN m^6^A RNA Enrichment (MeRIP) followed by dPCR using *circMbl*-specific probes. We find that m^6^A modification of *circMbl* is significantly elevated in tau transgenic flies compared to control (**Fig. 3G)**.

### m^6^A writers and readers modify *circMbl* formation and tau neurotoxicity

Having found that circRNAs are methylated to a higher degree in tau transgenic *Drosophila* and that *circMbl* is m^6^A-modified, we next asked if tau-induced elevation of m^6^A is mechanistically linked to an increase in RNA circularization. In *Drosophila*, m^6^A deposition on RNA is catalyzed by Mettl3 and Mettl14, while YTHDF1 detects and reads m^6^A to initiate downstream m^6^A-dependent RNA handling (Lence et al., 2017). To determine if genetic manipulation of RNA methylation writers and readers mediates *circMbl* abundance, we genetically reduced or overexpressed Mettl3, Mettl14, and YTHDF in neurons of tau^R406W^ transgenic *Drosophila*. Pan-neuronal RNAi-mediated knockdown of *Mettl3*, *Mettl14*, or *YTHDF* significantly reduced *circMbl*, suggesting that the tau-dependent increase of m^6^A regulates *circMbl* accumulation. Genetic overexpression of *Mettl3*, *Mettl14*, or human *YTHDF* (*hYTHDF*) did not alter *circMbl* levels in brains of tau transgenic *Drosophila* (**Fig. 4A**), suggesting that *circMbl* sites of potential m^6^A addition are already saturated in this model. Levels of transgenic tau protein were unchanged in response to genetic manipulation of m^6^A readers and writers (**Supplemental Fig. 2A, B**). As a control for the effects of genetic manipulation of m^6^A readers and writers on the expression of a normal, linear housekeeping gene, we analyzed levels of *β-tubulin* RNA in the context of *Mettl3*, *Mettl14*, and *YTHDF* knockdown and overexpression. *β-tubulin* transcript levels were unchanged as a consequence of genetic manipulation of RNA methylation machinery (**Fig. 4B**), suggesting that the effects of *Mettl3*, *Mettl14*, and *YTHDF* knockdown on *circMbl* are not a general consequence of widespread changes in transcript levels.

**Figure 4.**
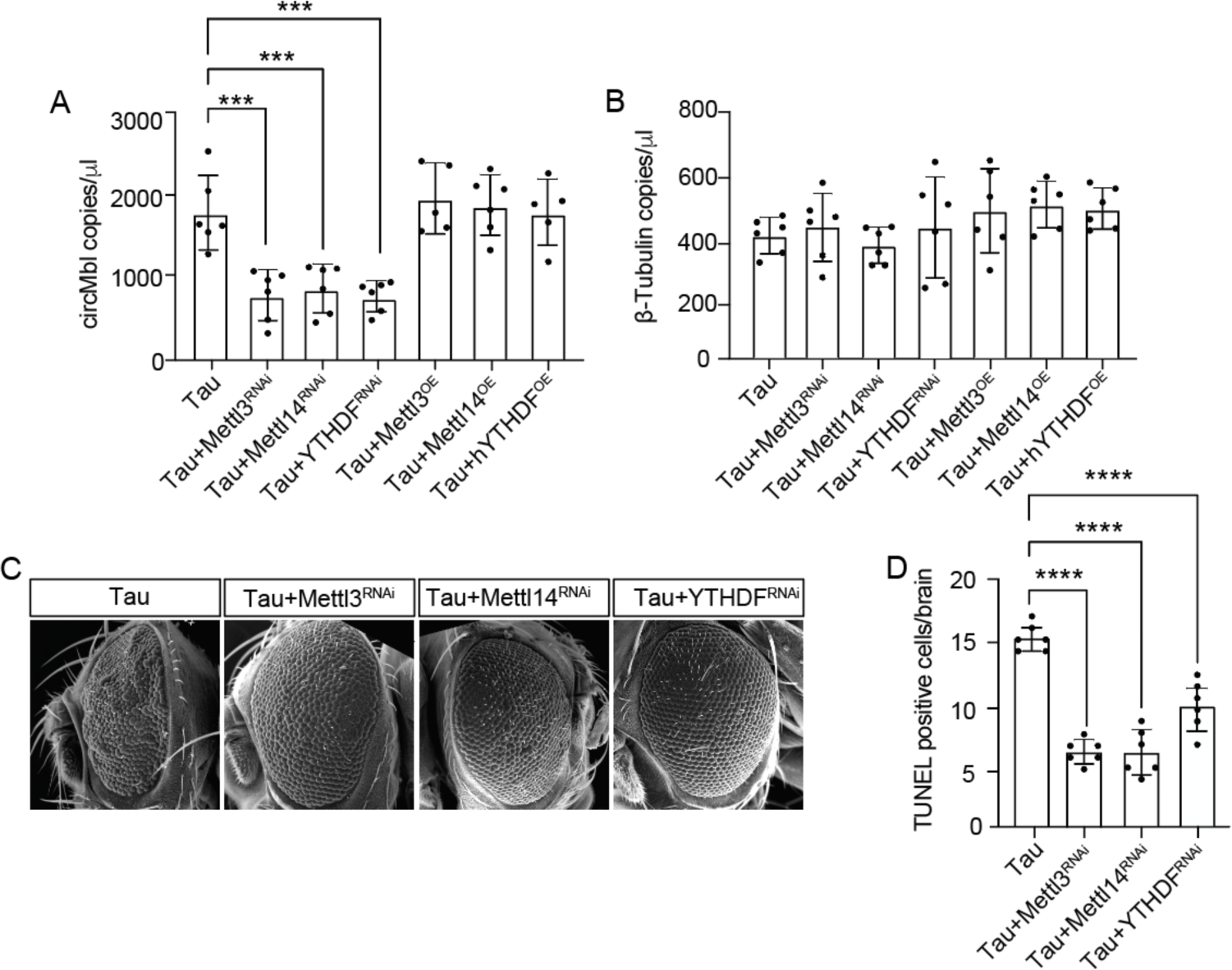
m^6^A writers and readers are involved in *circMbl* methylation. **A**. dPCR-based analysis of *circMbl* levels in *Drosophila* of the indicated genotype. **B**. dPCR analysis of *β-tubulin* levels in the indicated *Drosophila* genotypes. **C**. SEM analysis of the rough eye phenotype in *Drosophila* of the indicated genotype. **D**. Quantification of neurodegeneration based on TUNEL staining in *Drosophila* brains of the indicated genotype. n = 6 biological replicates per genotype. ***p,0.001, ****p<0.0001, ANOVA with a post-hoc Tukey test was used for multiple comparisons. Error bars = SEM. All analyses were performed at day 10 of adulthood.

Having found that m^6^A is elevated in tau transgenic *Drosophila* and that such elevation is a mechanistic driver of RNA circularization in this model, we next asked if tau-induced increase in RNA methylation causally mediates neurodegeneration. We find that panneuronal reduction of deposition and detection of RNA methylation factors suppresses the tau-induced rough eye phenotype (**Fig. 4C**), suggesting that tau-induced elevation of m^6^A is neurotoxic. To assess neurodegeneration in the brain directly, we quantified the number of TUNEL-positive cells in brains of tau transgenic *Drosophila* with and without RNAi-mediated depletion of *Mettl3*, *Mettl14*, and *YTHDF*. In line with our rough eye analyses, we find that genetic reduction of these factors suppresses tau-induced neurodegeneration (**Fig. 4D**).

### *CircMbl* and m^6^A associate with nuclear blebs in brains of tau transgenic *Drosophila*

Studies in *Drosophila,* iPSC-derived neurons, and human disease report the formation of nuclear invaginations and blebs in the context of tauopathy (Frost et al., 2016; Paonessa et al., 2019). As we have previously found that polyadenylated RNA and targets of nonsense-mediated RNA decay accumulate within nuclear invaginations and blebs in tau transgenic *Drosophila* (Cornelison et al., 2019; Zuniga et al., 2022), we visualized the relationship between *circMbl*, m^6^A, and nuclear pleomorphisms. Using transmission electron microscopy (TEM), we detect distinct structures within nuclear blebs in brains of tau transgenic *Drosophila*, the contents of which are unknown (**Fig. 5A**, **Supplemental Fig. 3**). We next performed FISH/IF to visualize *circMbl* with respect to nuclear pleomorphisms. In brains of tau transgenic *Drosophila,* but not controls, we find that *circMbl* associates with lamin-lined nuclear blebs (**Fig. 5B**). Similarly, we detect presence of m^6^A within lamin-positive nuclear blebs in brains of tau transgenic *Drosophila* (**Fig. 5C**). FISH/IF-based detection of *circMbl* and m^6^A reveals frequent colocalization of these factors in the perinuclear region (**Fig. 5D**). Taken together, these analyses suggest a potential relationship between circRNA, RNA methylation, and nuclear blebbing.

**Figure 5.**
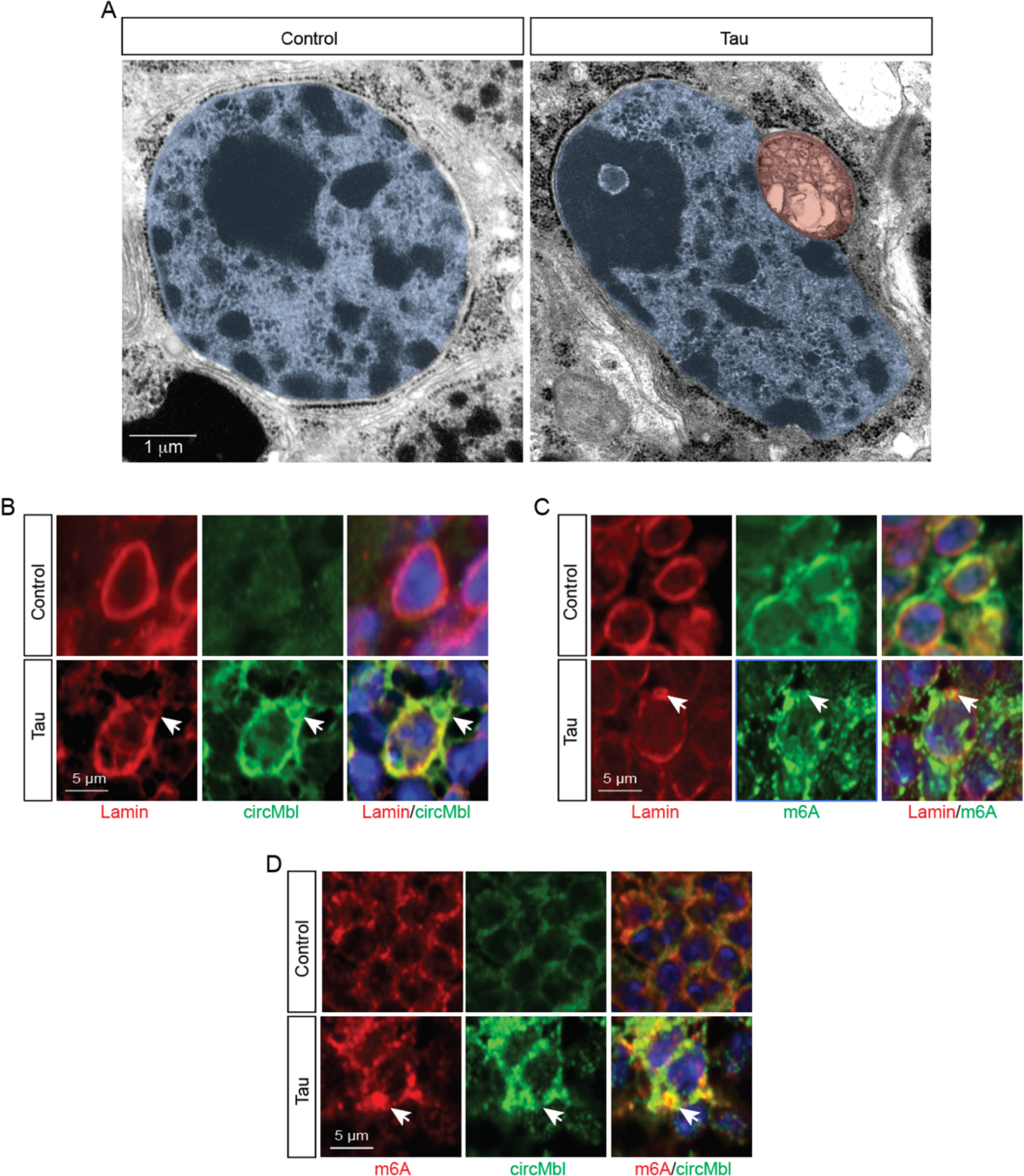
Association of *circRNA* and RNA methylation with tau-induced nuclear blebs in the *Drosophila* brain. **A**. Example of a nuclear bleb in the brain of tau transgenic *Drosophila* versus control visualized by TEM. The nucleus and nuclear bleb are artificially colored blue and red, respectively, for visualization. **B**. *CircMbl* associates with tau-induced nuclear blebs based on FISH/IF-based detection of *circMbl* and lamin in the *Drosophila* brain. Arrow indicates a lamin-lined nuclear bleb containing *circMbl*. **C**. m^6^A within a nuclear bleb of tau transgenic *Drosophila* based on visualization of m^6^A and lamin. Arrow indicates a lamin-lined nuclear bleb containing m^6^A. **D**. Accumulation of *circMbl* and m^6^A at the nuclear periphery in brains of tau transgenic *Drosophila*. Arrow indicates an example of colocalization. All flies were analyzed at day 10 of adulthood.

### Analyses of circRNA and m^6^A in iPSC-derived neurons and cerebral organoids carrying a ***MAPT* mutation**

We next investigated the incidence of circRNA formation and its relationship with RNA methylation in iPSC-derived neurons and 120-day-old cerebral organoids derived from a patient harboring the frontotemporal dementia-associated tau^IVS10+16^ *MAPT* mutation (Janssen et al., 2002) compared to its CRISPR-corrected isogenic control (Karch et al., 2019). We first utilized DCC to quantify circRNAs (**Supplemental Dataset 2**) in a publicly-available RNA-seq dataset from tau^IVS10+16/+^ iPSC-derived neurons compared to their CRISPR-corrected isogenic control (**Fig. 6A**). The human homolog of *Drosophila mbl, MBNL1,* was not detected in circular form in iPSC-derived neurons. In line with our findings in tau transgenic *Drosophila,* we find that overall levels of m^6^A are elevated in tau^IVS10+16^ iPSC-derived cerebral organoids aged to 120 days based on immunofluorescence (**Fig. 6B, C**). As has been reported for iPSC-derived neurons carrying other *MAPT* mutations (Paonessa et al., 2019), we also detect an increased incidence of nuclear envelope pleomorphism in 120-day-old tau^IVS10+16^ organoids based on immunofluorescence (**Fig. 6D, E**) and TEM (**Fig. 6F**, **Supplemental Fig. 4**).

**Figure 6.**
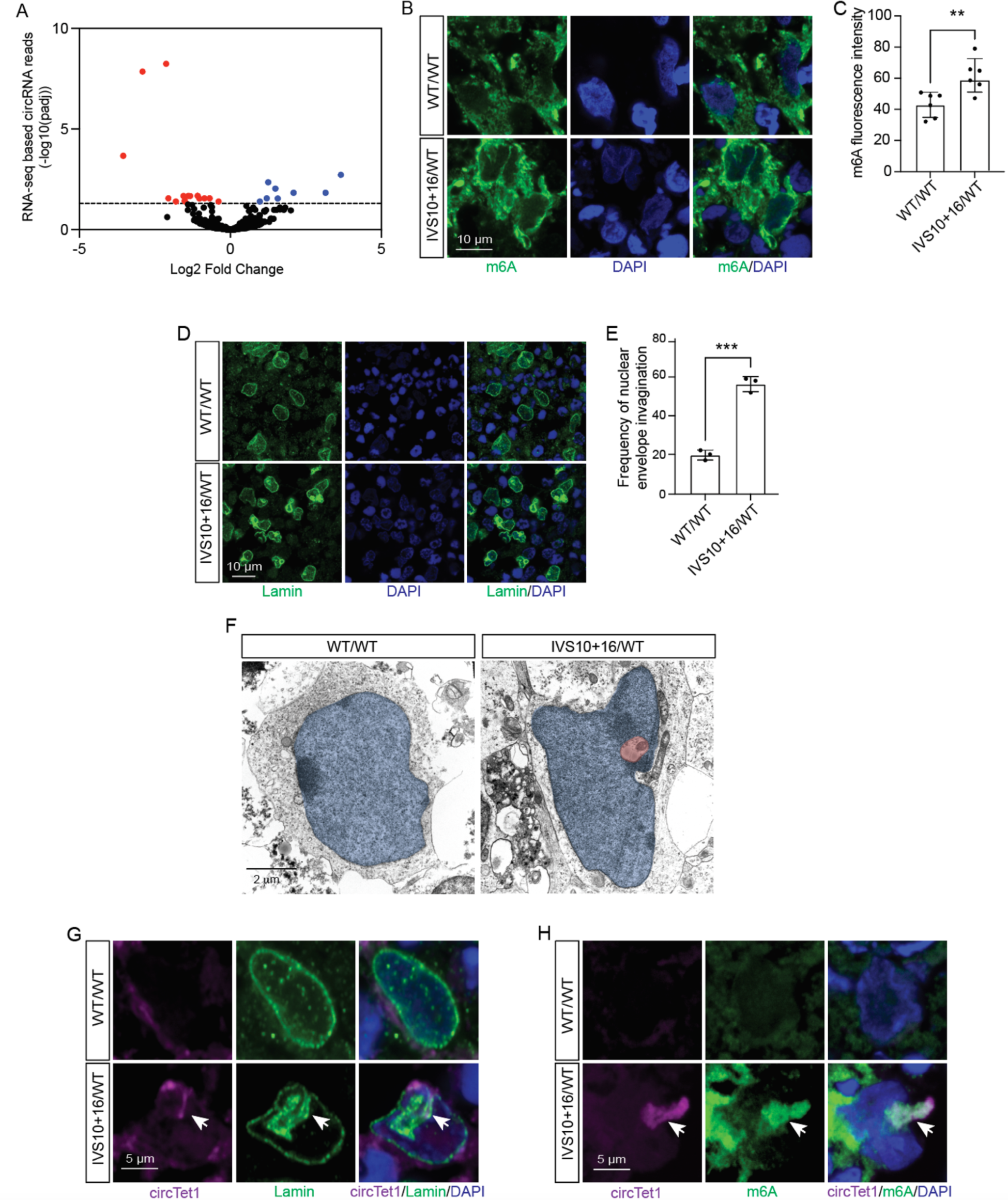
Differential circRNA expression, nuclear envelope invagination, and m^6^A modification in iPSC-derived neurons and cerebral organoids with *MAPT* mutation. **A.** RNA-seq based identification of differentially expressed circRNAs in tau^IVS10+16/+^ neurons compared to isogenic control. Dashed line indicates an adjusted p value of 0.05. **B**. Visualization of m^6^A in tau^IVS10+16/+^ iPSC-derived organoid and control based on immunofluorescence. **C**. Quantification of (B). **D**. Nuclear invagination based on lamin immunofluorescence in tau^IVS10+16/+^ iPSC-derived cerebral organoids and isogenic controls. **E**. Quantification of (D). **F**. TEM images showing large nuclear bleb with inclusion in tau^IVS10+16/+^ iPSC-derived cerebral organoid. The nucleus and inclusion are artificially colored blue and red, respectively, for visualization. FISH/IF-based detection of **G**. *circTet1* and Lamin B1 or **H.** m^6^A in human cerebral organoids. n = 6 replicates. *p<0.5, ***p,0.001, ***p<0.0001, unpaired t-test. Error bars = SEM. Brain organoids were aged to 120 days.

Having identified circRNAs that are differentially expressed in tau^IVS10+16^ iPSC-derived neurons, alongside an increase in m^6^A, we next visualized the relationship between circRNA, m^6^A, and nuclear pleomorphisms. We developed FISH probes to detect the back-splice junction of *circTet1*, a circRNA that was significantly enriched in tau^IVS10+16^ iPSC-derived neurons based on RNA-seq. FISH-based detection of *circTet1* confirms an overall increase of *circTet1* in tau^IVS10+16^ derived cerebral organoids, with clear presence in nuclear invaginations (**Fig. 6G and Supplemental Fig. 4A, B**). We also find that *circTet1* colocalizes with m^6^A in tau^IVS10+16^ derived cerebral organoids (**Fig. 6H**). In support of our mechanistic studies in *Drosophila*, our analyses of human *MAPT* mutation-carrying brain organoids suggests that tau dysfunction causes differential abundance of circRNAs, elevates RNA methylation, and drives nuclear envelope pleomorphisms that are associated with circRNA and m^6^A.

## DISCUSSION

With stable structure and long half-life in cerebrospinal fluid, circRNAs are a recently-proposed new hallmark of aging and potential biomarkers for neurodegenerative disorders (Gruner et al., 2016; Shao & Chen, 2016; Gokool et al., 2020). While circRNA dysregulation has been reported in age-associated neurological disorders including inflammatory neuropathy (Shao & Chen, 2016), Parkinson’s disease (Hanan et al., 2020) and Alzheimer’s disease (Dube et al., 2019), the association between pathogenic tau and circRNA biogenesis was previously unexplored. In the current study, we utilize a well-characterized *Drosophila* model of tauopathy and iPSC-derived neurons and cerebral organoids to discover a novel link between tau, m^6^A RNA methylation, circRNA biogenesis, nuclear pleomorphism and neurotoxicity.

We find that overall levels of circRNAs are elevated in tau transgenic *Drosophila* and that RNAi-mediated reduction of *circMbl* significantly suppresses tau neurotoxicity. While it is currently unknown how *circMbl* exerts toxicity in tau transgenic *Drosophila,* previous studies report that circRNAs can impact cellular function by competing with linear RNA splicing (Ashwal-Fluss et al., 2014), by serving as a sponge for complementary short and long RNAs and RNA binding proteins (Hansen et al., 2013; Memczak et al., 2013; Memczak et al., 2015), and by active translation into protein via cap-independent translation (Wang & Wang, 2015).

We became interested in m^6^A modification as a potential mechanism driving tau-induced circRNA formation based on previous studies reporting that differential m^6^A modification at specific sites is sufficient to alter the fate of a nascent RNA from its canonical splicing pattern to back-splicing and consequent circularization, and that the back-splicing rate of m^6^A-modified exons is significantly elevated compared to their unmethylated counterparts (Di Timoteo et al., 2020). The m^6^A modification occurs co-transcriptionally and is the most abundant and reversible RNA modification in eukaryotes (Desrosiers et al., 1974; X. Jiang et al., 2021). Writers, readers and erasers of m^6^A modification dynamically edit RNAs and affect RNA splicing, export, translation and clearance (X. Jiang et al., 2021). We find that transgenic expression of human tau causes an overall increase of m^6^A in the brain, that circRNA is m^6^A modified to a greater extent in brains of tau transgenic *Drosophila*, and that panneuronal RNAi-mediated depletion of m^6^A readers and writers (*Mettl3, Mettl14 and YTHDF*) reduce *circMbl* content in tau transgenic *Drosophila*.

As a previous study utilizing transgenic expression of *R406W* mutant tau in the *Drosophila* eye reported exacerbation of the tau-induced rough eye phenotype upon RNAi-mediated knockdown of *Mettl3, Mettl14 and YTHDF* (Shafik et al., 2021), we were surprised to find robust suppression of the tau-induced rough eye and a decrease in TUNEL-positive cells per brain in response to panneuronal knockdown of these m^6^A regulators in *Drosophila* with panneuronal expression of tau^R406W^. We would not expect such discrepancy between our respective analyses despite minor differences in our approach – use of a panneuronal driver in our study rather than eye-specific transgene expression, for example, or our quantification of neurotoxicity via TUNEL rather than relying on the rough eye phenotype alone. Our findings do nevertheless align with recent work reporting an overall increase in m^6^A in brains of tau transgenic mice and with increasing Braak stage in human Alzheimer’s disease (L. Jiang et al., 2021). In this study, multimeric forms of tau were found to indirectly associate with m^6^A-modified RNA via direct binding of tau to HNRNPA2B1, an m^6^A reader. In line with our current work, the association between tau and HNRNPA2B1 (and, indirectly, m^6^A modified RNA) was found to be neurotoxic, as siRNA-mediated knockdown of HNRNPA2B1 reduced apoptotic cell death as measured by cleaved caspase-3 in cultured neurons with induced tau aggregation and in young PS19 tau transgenic mice injected with multimeric forms of tau. The m^6^A writer METTL3 is significantly elevated in the insoluble fraction of total brain lysates from postmortem human Alzheimer’s disease hippocampus, further implicating RNA methylation machinery in human tauopathy (Huang et al., 2020). While both circRNA and m^6^A dysregulation have thus both been previously implicated in human Alzheimer’s disease, our study is the first to identify m^6^A dysregulation as a mechanistic driver of tau-induced differential circRNA expression.

CircRNAs can be classified into three main categories: exonic circRNAs (EcircRNA), which comprise exonic regions; exonic-intronic circRNAs (EIcircRNA), consisting of both exonic and intronic regions; and intronic circRNAs (IcircRNA), composed solely of intronic elements (Meng et al., 2017). Cytoplasmic circRNAs, predominantly EcircRNAs, possess cytoplasmic localization signals and are thus exported into the cytosol, whereas IcircRNAs remain confined to the nucleus. EcircRNAs function as microRNA sponges and can undergo translation, while IcircRNAs play a role in regulating gene transcription by modifying splicing patterns (Zhou et al., 2021). CircMbl is classified as an EcircRNA as it is formed by the back-splicing of exon 2. Accordingly, we detect very low levels of circMbl in the nucleus, suggesting that it is actively exported.

Given the differential expression of circRNAs in tau transgenic *Drosophila*, *MAPT* mutant iPSC-derived neurons, and in human Alzheimer’s disease brain (Dube et al., 2019), alongside our previous discovery that polyA RNA and error-containing RNAs accumulate within tau-induced nuclear envelope invaginations and blebs (Cornelison et al., 2019), we investigated the association between nuclear pleomorphisms and circRNAs in models of tauopathy. We find that *circMbl* and *circTet1* are significantly enriched within nuclear blebs of tau transgenic *Drosophila* and *MAPT* mutant iPSC-derived cerebral organoids, respectively. While nuclear pore complexes embedded in the nuclear envelope facilitate the transport of most RNAs and proteins between the nucleus and cytosol, certain larger macromolecular complexes exit the nucleus through nuclear envelope budding (Speese et al., 2012; Tanabe et al., 2016). During nuclear budding, large inclusions such as RNA binding complexes are encapsulated within the inner nuclear membrane and extend into the perinuclear space before being released into the cytosol (Speese et al., 2012). Based on electron microscopy, we detect what appear to be nuclear buds in brains of tau transgenic *Drosophila* and in *MAPT* mutant cerebral organoids. While we do not currently know if tau-induced nuclear budding or circRNAs cause these structures, we speculate that circRNA-mediated sequestration of RNAs and RNA binding proteins leads to the formation of large macromolecules that induce nuclear budding as a nuclear export mechanism.

Muscleblind has previously been implicated in various neurological disorders associated with nucleotide repeat expansions, including myotonic dystrophy (Kanadia et al., 2003; de Haro et al., 2006) spinocerebellar ataxia (Li et al., 2008; Daughters et al., 2009), fragile X syndrome (Sellier et al., 2010), and Huntington’s disease (Rudnicki et al., 2007). Recent work also points toward muscleblind as a mediator of toxicity in amyotrophic lateral sclerosis (ALS) associated with mutation of the splicing factor fused in sarcoma (*FUS*) (Casci et al., 2019). Previous genetic screening of *Drosophila* models of FUS (Casci et al., 2019) and ataxin 3 polyglutamine expansion (Li et al., 2008) identified mbl as a modifier of neurotoxicity; both studies report suppression of toxicity upon reduced mbl levels/function, pointing toward a gain-of-function mechanism of mbl-associated toxicity. Similarly, knockdown of the human or mouse orthologues of mbl, muscleblind-like splicing regulator 1 (*MBNL1* (Hs) and *Mbnl1* (Mm)), significantly reduce the burden of FUS-associated stress granules in HEK293T cells and rat primary cortical neurons (Casci et al., 2019). Studies in myotonic dystrophy, however, point toward loss of MBNL1 function as a disease mechanism. MBNL1 deposits within nuclear inclusions in this disorder due to its binding to CUG repeat expansions in mRNA transcripts of *dystrophia myotonica protein kinase*, thus reducing MBNL1 function as a splicing factor (Miller et al., 2000). Our work adds tauopathy to the list of neurological disorders that may be associated with mbl dysfunction and point toward increased m^6^A methylation and circularization of the *mbl* transcript as a mechanism driving disease. While it is currently unknown if circularization of muscleblind transcripts are involved in FUS-associated neurotoxicity or in various neurological disorders involving nucleotide repeat expansion, our investigations into *circMbl* and tau neurotoxicity provide strong rationale for investigating *MBNL1* transcript circularization in these disorders, as well as in the adult human brain affected by tauopathy.

Taken together, we have discovered a toxic link between tau, elevated levels of RNA methylation, and formation of circRNAs. While the current study utilizes two different frontotemporal dementia-associated mutations of human tau, the presence of nuclear envelope pleomorphisms alongside the dysregulation of circRNA and RNA methylation previously reported in human Alzheimer’s disease and other models of tauopathy (Frost et al., 2016; Cornelison et al., 2019; Dube et al., 2019; L. Jiang et al., 2021) suggests that our findings are likely not a restricted to the *R406W* or *IVS10+16 MAPT* mutations. In future work, it will be of interest to further define the role of m^6^A-mediated circRNA formation, nuclear export, and translation in the setting of sporadic primary and secondary tauopathy.

## MATERIALS AND METHODS

### Drosophila genetics

*Drosophila* crosses and aging were performed at 25 °C with a 12-hour light/dark cycle on a standard diet (Bloomington formulation). Full *Drosophila* genotypes and sources are listed in **Supplemental Table 1**. Panneuronal expression of transgenes in *Drosophila* utilized the GAL4/UAS system with the *elav* promoter driving GAL4 expression. All analyses utilized an equal number of male and female flies.

### Cell culture

Human iPSC lines were obtained from the Tau Consortium cell line collection (https://www.neuralsci.org/tau) (Karch et al., 2019). All iPSC lines were maintained in complete mTeSR1 (#85850; Stem Cell Technology) growth medium on Corning® Matrigel® hESC-Qualified Matrix (#354277; Corning) and passaged every 3-5 days using ReLeSR™ (#05872; Stem Cell Technology).

Cortical organoids were generated as previously described (Paşca et al., 2015) with minor modifications. iPSC cultures at ~80% confluency were detached from plates using ReLeSR™, the cell pellet was mixed gently with STEMdiff™ Neural Induction Medium (NIM)+1µl/ml ROCK inhibitor Y-27632 (#72302; Stem Cell Technology) and plated at 3×10^6^ cell density per well into Nunclon Sphera 96 U bottom plates (#174929; ThermoFisher Scientific) and incubated at 37 °C with 5% CO_2_. From days 2-6, fresh NIM media supplemented with dorsomorphin (#P5499; Sigma-Aldrich) and SB-431542 (#1614; Tocris) were added to cells daily. From days 7-24, spheroids were fed every other day with STEMdiff™ Neural Progenitor Medium (#05833; Stem Cell Technology). After day 23, media was replaced every other day with neuronal differentiation media prepared based on the Stem Cell Technology protocol: 10 ml of the BrainPhys™ Neuronal Medium (#05790; Stem Cell Technology) supplemented with 200 µl of NeuroCultTM SM1 Neuronal Supplement (#05711; Stem Cell Technology), 200 µl of N2 Supplement-A (#07152; Stem Cell Technology), 20 ng/ml Human Recombinant BDNF (#78005; Stem Cell Technology), 20 ng/ml Human Recombinant GDNF (#78058; Stem Cell Technology), 1 mM Dibutyryl-cAMP (#73882; Stem Cell Technology) and 200 nM Ascorbic Acid (#72132; Stem Cell Technology). From day 43 to 120, spheroids were fed every four days with BrainPhys™ Neuronal Medium.

### Bioinformatic analyses

*Drosophila.* Total RNA sequencing data from 10-day old *Drosophila* transgenic tau^R406W^ vs control heads (GSE152278) were aligned to the *Drosophila melanogaster* genome (Version 6.50) with STAR v2.7.1a (--chimSegmentMin 15 and --chimJunctionOverhangMin 15). CircRNAs were identified and quantified with DCC v0.50 (Cheng et al., 2016) with Flybase gene annotations (v6.50), RepeatMasker and UCSC simple repeat filtered annotation file using default parameters. Detected circRNAs and other transcripts were then tested for differential expression using DESeq2 v1.34. Differentially expressed circRNAs with an adjusted p value less than 0.05 were considered statistically significant. Sequencing data are provided in **Supplemental Dataset 1**.

*iPSC-derived neurons*. Fastq files were aligned to the GRCh38 genome with STAR v2.7.1a (-- chimSegmentMin 15 and --chimJunctionOverhangMin 15). CircRNAs were identified and quantified with DCC v0.50 (Cheng et al., 2016) with RepeatMasker and UCSC simple repeat filtered annotation file using default parameters. Detected circRNAs and transcripts were then tested for differential expression using DESeq2 v1.34. Differentially expressed circRNAs with an FDR less than 0.05 were considered statistically significant. Sequencing data are provided in **Supplemental Dataset 2**.

### dPCR

Absolute RNA levels of *mbl*, *circMbl* and *tubulin* were assessed using the Applied Biosystems QuantStudio 3D dPCR system. For RNA extraction, six fly heads (three males and three females) were homogenized in TRIzol (Invitrogen) and RNA was extracted according to the manufacturer’s protocol. RNA concentrations were measured using a Nanodrop8000 spectrophotometer (ThermoFisher Scientific). Equal quantities of RNA were added to a reverse transcription reaction (cDNA Reverse Transcription Kit, Applied Biosystems). Resulting cDNA was loaded into a dPCR chip, sealed, and amplified prior to calculating the absolute RNA concentration in copies/μL. Probes for linear *mbl* and *tubulin* mRNA were predesigned by ThermoFisher Scientific. The *circMbl* probe was designed to recognize the junction between the 5’ and 3’ ends of the RNA using the Custom TaqMan^®^ Assay Design Tool (**Supplemental Table 3)**.

### Western blotting

One *Drosophila* head per lane was homogenized in 2X Laemmeli buffer (Bio-Rad) and boiled for 10 minutes. Protein was separated via 10% SDS-PAGE and transferred to nitrocellulose membranes. Membranes were incubated overnight at 4 °C with primary antibodies (**Supplemental Table 4**). After washing with PBS+0.1% Tween (PBSTw), membranes were incubated with HRP-conjugated secondary antibodies for two hours at room temperature and developed with Clarity^TM^ Western ECL substrate. Images were acquired using the ChemiDoc^TM^ imaging system (Bio-Rad), after which FIJI was used to quantify signal intensity.

### Fluorescence *in situ* hybridization/immunofluorescence (FISH-IF)

Frozen OCT-embedded *Drosophila* heads were sectioned at 10 μm, placed on glass slides, and fixed with 4% PFA for 10 minutes. Tissue was incubated in pre-hybridization buffer (2X SSC, 10% dextran sulfate, 20 mM ribonucleoside vanadyl complex (RVC), 20% formamide) for 15 minutes at room temperature. Digoxigenin (DIG)-labeled *circMbl* and Quasar®670-labeled *circTet1* probes (**Supplemental Table 5**) were mixed with hybridization buffer (4X SSC, 20% dextran sulfate, 40 mM RVC) at 2 ng/µl concentration and incubated for three hours at 37 °C followed by three 10-minute washes in 2X SSC at 37 °C. Sections were then incubated with anti-DIG (Invitrogen) with 20 mM of RVC for one hour at 37 °C and washed briefly with PBS + 0.2% Triton-X 100 (PBSTr). Preparations were fixed in 4% paraformaldehyde for 10 minutes and washed three times in PBSTr for 10 minutes each followed by overnight incubation with antibodies detecting lamin (lamin R-836 (Osouda et al., 2005), and m6A at 4 °C. Samples were then incubated with AlexaFluor^™^-conjugated secondary antibodies for two hours at room temperature and briefly rinsed with PBSTr and mounted with DAPI (Southern Biotech).

For visualization of lamin and m^6^A in *Drosophila* brains, we utilized formalin fixed, paraffin embedded *Drosophila* heads sectioned at 4 μm. Sections were rehydrated and then subjected to antigen retrieval via microwaving in 1 mM of sodium citrate for 20 minutes. Slides were blocked with 2% BSA in PBSTr and incubated with primary antibodies overnight at 4 °C. Slides were washed again prior to incubation with AlexaFluor-conjugated secondary antibodies for two hours at room temperature. Samples were visualized using a Zeiss confocal microscope.

### Electron microscopy

Fly heads or iPSC-derived organoids were fixed in phosphate buffered 4% formaldehyde and 1% glutaraldehyde for at least two hours followed by osmium tetroxide treatment for 30 minutes. The samples were then dehydrated through a graded series of ethanol. For Scanning Electron Microscopy (SEM), samples were treated with hexamethyldisilazane and air dried before sputter coated with gold/palladium. For Transmission Electron Microscopy (TEM), each piece was embedded with resin in a flat mold for correct orientation and then polymerized in an 85 °C oven overnight. Images were captured using JEOL JSM 6610LV and JEOL 1400 for SEM and TEM, respectively.

### *Drosophila* locomotor assay

The *Drosophila* locomotor assay was performed as previously described (Frost et al., 2016). Briefly, *Drosophila* were individually transferred into a new vial at day nine of adulthood. On day 10, vials were placed on a grided surface, and the number of centimeters walked in 30 seconds was quantified. 18 flies per genotype were analyzed. Investigators were blinding to genotype.

### TUNEL

Formalin-fixed, paraffin embedded *Drosophila* heads were sectioned at 4 μm. TdT Colorimetric FragEL DNA Fragmentation Detection Kit (QIA33 TdT FragEL; Calbiochem) was used for TUNEL staining. Diaminobenzidine was used for secondary detection of biotin-labeled deoxynucleotides at exposed ends of DNA fragments per the manufacturer’s recommendation. Brightfield microscopy was used to quantify TUNEL-positive neurons throughout the entire fly brain. Investigators were blinded to genotype.

### Dot blot

RNA was extracted as described above and was serially diluted to 25, 12.5, 6.25 ng/µl. Samples were loaded on a positively charged nylon membrane (Bright Star-Plus; Invitrogen) and UV crosslinked using Stratagene Stratalinker 1800. Membranes were washed briefly with PBSTw and blocked in PBSTw plus 2% dry milk for 30 minutes followed by overnight incubation with the m^6^A antibody at 4 °C. The membrane was then incubated with an HRP-conjugated secondary antibody for two hours at room temperature and developed with Clarity^TM^ Westrern ECL substrate (Bio-Rad). Images were acquired using the ChemiDoc^TM^ imaging system (Bio-Rad), after which FIJI was used to quantify signal intensity.

### Me-RIP

RNA was extracted as described above from 80 *Drosophila* heads (40 females and 40 males). m^6^A enrichment was performed using the EpiQuik CUT&RUN m6A RNA Enrichment (MeRIP) Kit (#P-9018-24, EPIGENTEK) based on the manufacturer protocol and 20 µg (m6A enriched) or 200 ng (input) of RNA per sample. m6A enriched and input RNAs were eluted in nuclease free water for further cDNA preparation and dPCR analysis.

### RNaseR treatment

RNA was extracted from 12 *Drosophila* heads as described above. 2 µg of RNA per sample was digested using one unit of RNase R and 2 µl of 10X RNase R buffer (#RNR07250, Bioresearch Technologies) in a 20 µl reaction at 37 °C for 10 minutes. The resulting circRNA was then subjected to analysis via dot blot as described above.

### Statistical analyses

We considered *P* < 0.05 significant based on a student’s t-test and ANOVA followed by Tukey’s test when making single or multiple comparisons, respectively. Unless otherwise noted, statistical analyses were performed using GraphPad Prism 9.1. Based on power analysis and our previous studies, we used n = 6 biological replicates per condition for immunostaining, FISH/IF and Western blotting, n = 12 for digital PCR, and n = 18 for locomotor activity. When possible, samples were randomized and visual scoring was performed blindly for all immunofluorescence, TUNEL and locomotor assays.

## Acknowledgements

We thank Drs. Sebastian Kadener and Nagarjuna Pamudurti, Brandeis University, for providing the *circMbl*^KD^ *Drosophila* stock and Dr. Mel Feany for providing UAS-tau^R406W^ *Drosophila*. All other *Drosophila* stocks were obtained from the Bloomington Drosophila Stock Center (NIH P40OD018537). The actin antibody (JLA20) developed by Lin, J.J.-C (Lin, 1981) was obtained from the Developmental Studies Hybridoma Bank, created by the NICHD of the NIH and maintained at The University of Iowa, Department of Biology, Iowa City, IA 52242.

## Contributions

The study was conceptualized by FA and BF. FA, JD, EG and MM performed experiments. FA and PR completed data analysis. BF and FA participated in study design, figure and manuscript preparation. MM and CK provided RNA-seq data for iPSC-derived neurons prior to publication and deposition of this data in the Synapse repository.

## SUPPLEMENTAL INFORMATION

### Supplemental Figures

**Supplemental Figure 1.**
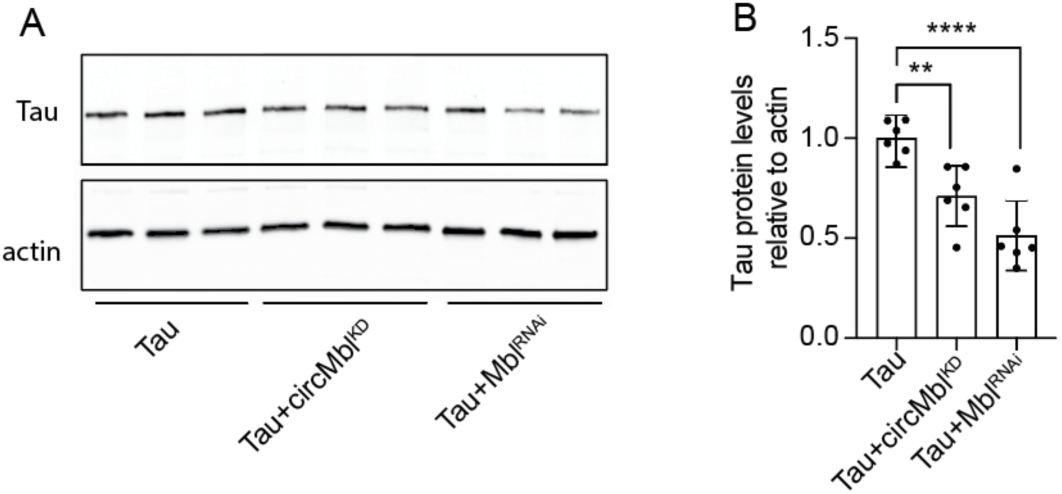
Tau protein levels in tau^R406W^ transgenic *Drosophila* with panneuronal reduction of *circMbl* or *mbl*. **A**. Tau protein levels in total head lysates from tau^R406W^, tau+*circMbl*^KD^ and tau+*Mbl*^RNAi^ *Drosophila.* **B**. Quantification of (C). n = 6 biological replicates per genotype. \*\**p* < 0.01, \*\*\*\**p* < 0.0001, ANOVA with a post-hoc Tukey test was used for multiple comparisons. All assays were performed at day 10 of adulthood. Error bars = SEM.

**Supplemental Figure 2.**
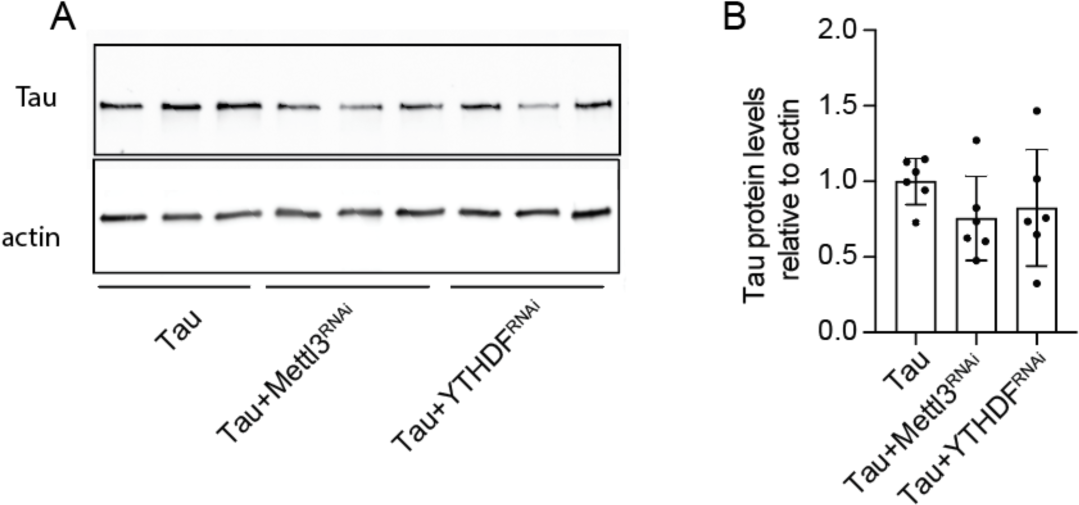
Tau protein levels in tau^R406W^ transgenic *Drosophila* with panneuronal reduction of *Mettl3* and *YTHDF*. **A**. Tau protein levels in total head lysates from tau^R406W^, tau+*Mettl3l*^RNAi^ and tau+*YTHDF^RNAi^ Drosophila.* **B**. Quantification of (A). *n* = 6 biological replicates per genotype. ANOVA with a post-hoc Tukey test was used for multiple comparisons. All assays were performed at day 10 of adulthood. Error bars = SEM.

**Supplemental Figure 3.**
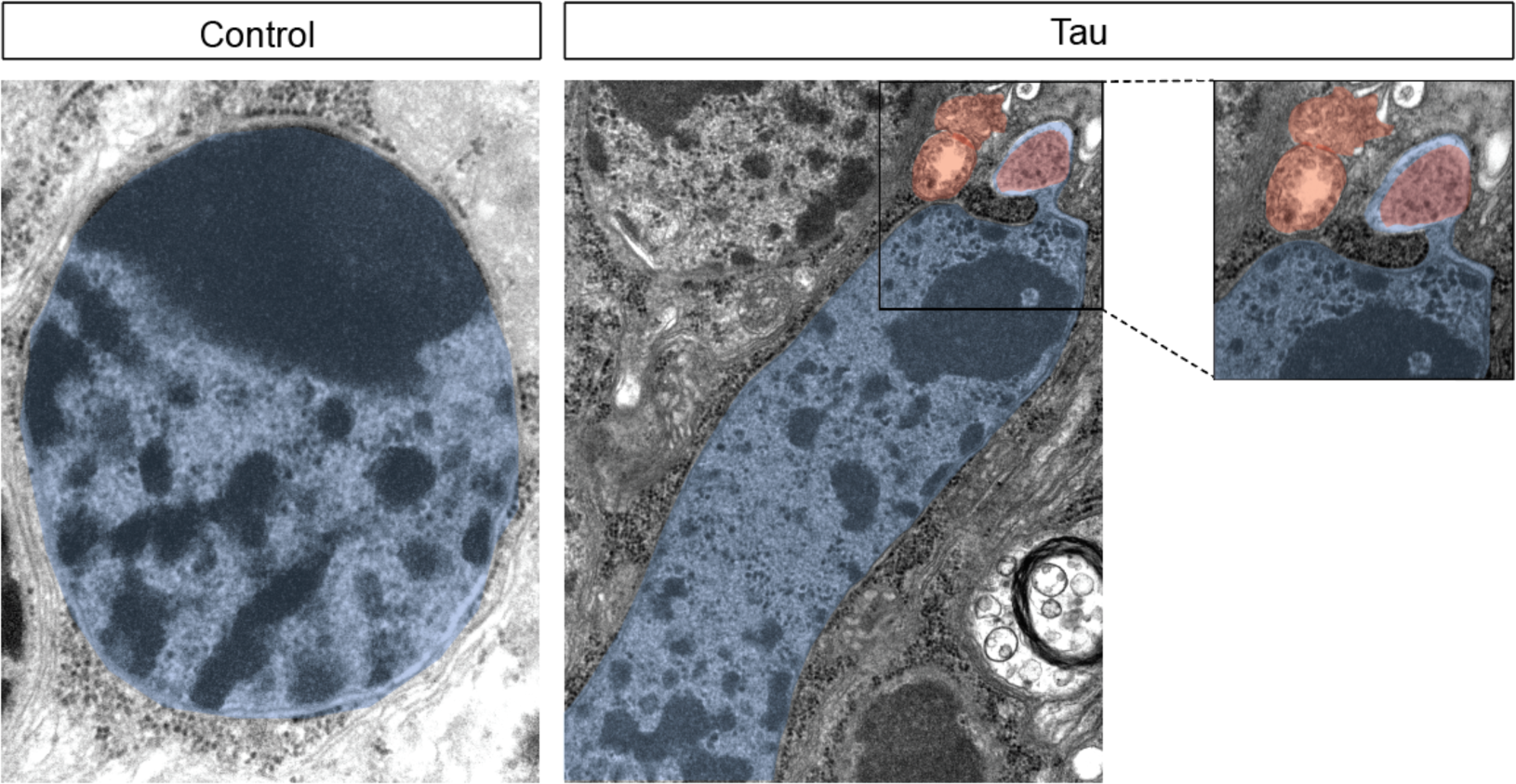
Nuclear bleb formation increases in neurons of tau transgenic *Drosophila*. A nuclear bleb containing a large inclusion in the brain of tau transgenic *Drosophila* versus control visualized by TEM showed in upper panel. Lower panel represents a protruding bleb in tau transgenic brain. The nucleus and inclusion are artificially colored blue and red, respectively, for visualization. All flies were 10 days old.

**Supplemental Figure 4.**
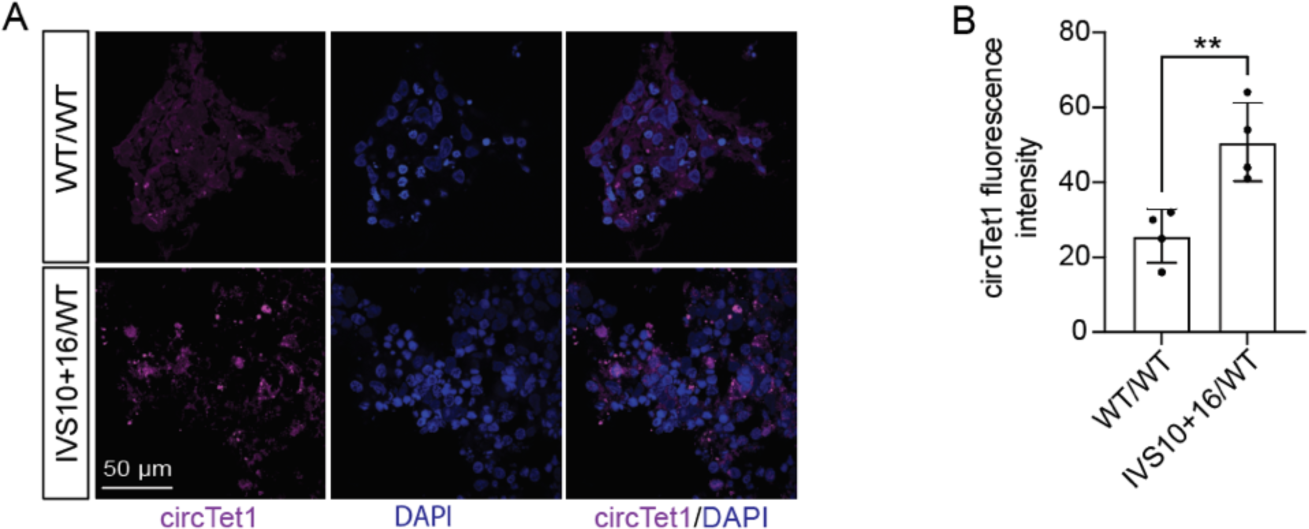
Elevated levels of *circTet1* in cortical brain organoids with *MAPT* mutation. **A.** Visualization of *circTet1* in tau^IVS10+16/+^ iPSC-derived cerebral organoid vs. isogenic control based on FISH. **B.** Quantification of *circTet1* levels in (A). **p<0.01, unpaired t-test.

### Supplemental Tables

**Supplemental Table 1.** *Drosophila* genotypes and sources.

| Name | Full Genotype | Stock number/source |
| --- | --- | --- |
| Control | elav-GAL4/+;+/+;+/+ | Elav-GAL4: BDSC 458 |
| Tau | elav-GAL4/+;+/+;UAS-Tau<sup>R406W</sup>/+ | Tau<sup>R406W</sup>: Mel Feany |
| Tau+circMbl<sup>KD</sup> | elav-GAL4/+;circMbl<sup>KD</sup>/+;UAS-Tau<sup>R406W</sup>/+ | circMbl<sup>KD</sup>: Sebastian Kadener |
| Tau+Mbl<sup>RNAi</sup> | elav-GAL4/+;+/+;UAS-Tau<sup>R406W</sup>/UAS-Mbl<sup>RNAi</sup> | Mbl<sup>RNAi</sup>: BDSC 29585 |
| Tau+YTHDF<sup>RNAi</sup> | elav-GAL4/+;UAS-YTHDF<sup>RNAi</sup>/+;UAS-Tau<sup>R406W</sup>/+ | YTHDF<sup>RNAi</sup>: BDSC 55151 |
| Tau+Mettl3<sup>RNAi</sup> | elav-GAL4/+;+/+;UAS-Tau<sup>R406W</sup>/UAS-Mettl3<sup>RNAi</sup> | Mettl3<sup>RNAi</sup>: BDSC 30448 |
| Tau+Mettl14<sup>OE</sup> | elav-GAL4/+;UAS-Mettl1<sup>OE</sup>/+;UAS-Tau<sup>R406W</sup>/+ | Mettl14<sup>OE</sup>: BDSC 479407 |
| Tau+Mettl3<sup>OE</sup> | elav-GAL4/+;+/+;UAS-Tau<sup>R406W</sup>/UAS-Mettl3<sup>OE</sup> | Mettl3<sup>OE</sup>: BDSC 86295 |
| Tau+hYTHDF<sup>OE</sup> | elav-GAL4/+;UAS-hYTHDF<sup>OE</sup>/+;UAS-Tau<sup>R406W</sup>/+ | hYTHDF<sup>OE</sup>: BDSC 91178 |

**Supplemental Table 2.**
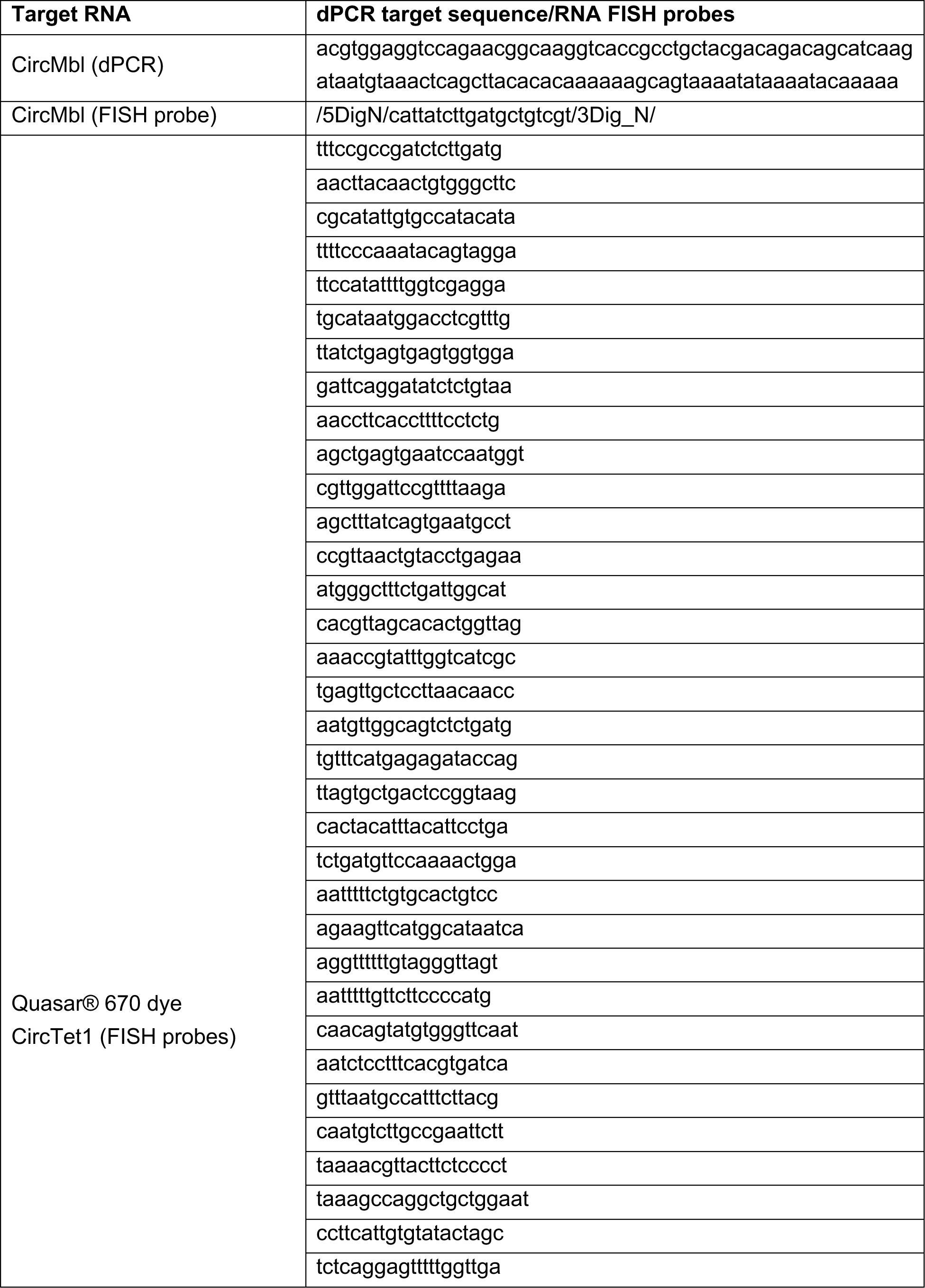
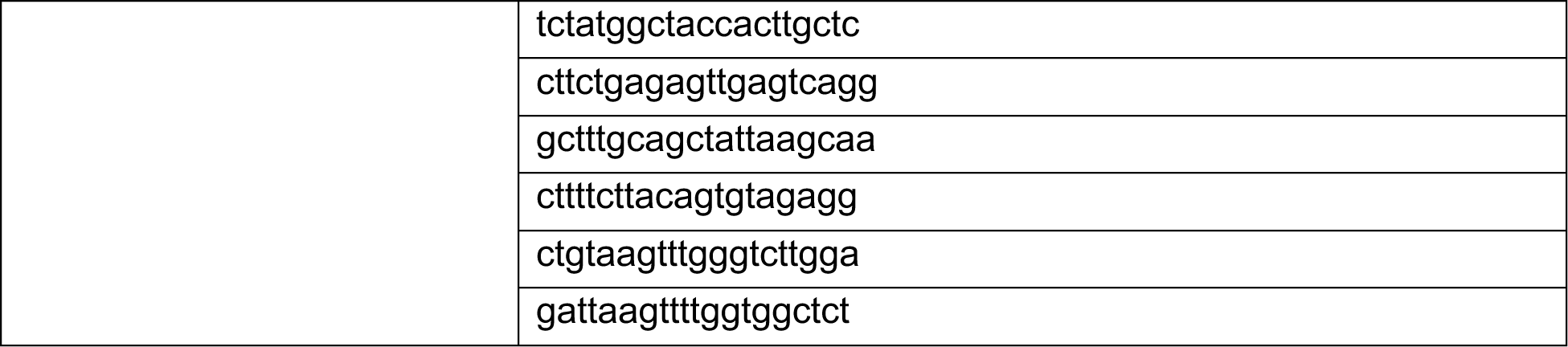
dPCR target sequence and RNA FISH probes.

**Supplemental Table 3.** List of antibodies, concentrations and sources.

| **Antibody** | **Western Blot** | **Immunofluorescence** | **Dot Blot** | **Source** |
| --- | --- | --- | --- | --- |
| Actin | 1:1000 |  |  | DSHB# JLA29 |
| Lamin R-836 |  | 1:200 |  | Paul Fisher |
| Lamin B1 |  | 1:100 |  | Abcam 16048 |
| m6A |  | 1:1,000 | 1:1,000 | Abcam 151230 |
| Digoxigenin |  | 1:100 |  | Invitrogen 700772 |
| Tau | 1:2,000,000 |  |  | Dako A0024 |
| Alexa 555 |  | 1:200 |  | Invitrogen A21424 |
| Alexa 488 |  | 1:200 |  | Invitrogen A11008 |
| HRP | 1:20,000 |  |  | Southern Biotech 1010-5 |
| HRP | 1:20,000 |  |  | Southern Biotech 3010-5 |

## REFERENCES

Apicco, D. J., Zhang, C., Maziuk, B., Jiang, L., Ballance, H. I., Boudeau, S., … Wolozin, B. (2019). Dysregulation of RNA Splicing in Tauopathies. Cell Rep, 29(13), 4377–4388 e4374. 10.1016/j.celrep.2019.11.093

Ashwal-Fluss, R., Meyer, M., Pamudurti, N. R., Ivanov, A., Bartok, O., Hanan, M., … Kadener, S. (2014). circRNA biogenesis competes with pre-mRNA splicing. Mol Cell, 56(1), 55–66. 10.1016/j.molcel.2014.08.019

Bardai, F. H., Wang, L., Mutreja, Y., Yenjerla, M., Gamblin, T. C., & Feany, M. B. (2018). A Conserved Cytoskeletal Signaling Cascade Mediates Neurotoxicity of FTDP-17 Tau Mutations. J Neurosci, 38(1), 108–119. 10.1523/JNEUROSCI.1550-17.2017

Braak, H., Thal, D. R., Ghebremedhin, E., & Del Tredici, K. (2011). Stages of the pathologic process in Alzheimer disease: age categories from 1 to 100 years. J Neuropathol Exp Neurol, 70(11), 960–969. 10.1097/NEN.0b013e318232a379

Casci, I., Krishnamurthy, K., Kour, S., Tripathy, V., Ramesh, N., Anderson, E. N., … Pandey, U. B. (2019). Muscleblind acts as a modifier of FUS toxicity by modulating stress granule dynamics and SMN localization. Nat Commun, 10(1), 5583. 10.1038/s41467-019-13383-z

Cheng, J., Metge, F., & Dieterich, C. (2016). Specific identification and quantification of circular RNAs from sequencing data. Bioinformatics, 32(7), 1094–1096. 10.1093/bioinformatics/btv656

Cornelison, G. L., Levy, S. A., Jenson, T., & Frost, B. (2019). Tau-induced nuclear envelope invagination causes a toxic accumulation of mRNA in Drosophila. Aging Cell, 18(1), e12847. 10.1111/acel.12847

Daughters, R. S., Tuttle, D. L., Gao, W., Ikeda, Y., Moseley, M. L., Ebner, T. J., … Ranum, L. P. (2009). RNA gain-of-function in spinocerebellar ataxia type 8. PLoS Genet, 5(8), e1000600. 10.1371/journal.pgen.1000600

de Haro, M., Al-Ramahi, I., De Gouyon, B., Ukani, L., Rosa, A., Faustino, N. A., … Botas, J. (2006). MBNL1 and CUGBP1 modify expanded CUG-induced toxicity in a Drosophila model of myotonic dystrophy type 1. Hum Mol Genet, 15(13), 2138–2145. 10.1093/hmg/ddl137

Desrosiers, R., Friderici, K., & Rottman, F. (1974). Identification of methylated nucleosides in messenger RNA from Novikoff hepatoma cells. Proc Natl Acad Sci U S A, 71(10), 3971–3975. 10.1073/pnas.71.10.3971

Di Timoteo, G., Dattilo, D., Centron-Broco, A., Colantoni, A., Guarnacci, M., Rossi, F., … Bozzoni, I. (2020). Modulation of circRNA Metabolism by m(6)A Modification. Cell Rep, 31(6), 107641. 10.1016/j.celrep.2020.107641

Dias-Santagata, D., Fulga, T. A., Duttaroy, A., & Feany, M. B. (2007). Oxidative stress mediates tau-induced neurodegeneration in Drosophila. J Clin Invest, 117(1), 236–245. 10.1172/JCI28769

Dube, U., Del-Aguila, J. L., Li, Z., Budde, J. P., Jiang, S., Hsu, S., … Cruchaga, C. (2019). An atlas of cortical circular RNA expression in Alzheimer disease brains demonstrates clinical and pathological associations. Nat Neurosci, 22(11), 1903–1912. 10.1038/s41593-019-0501-5

Dubin, D. T., & Taylor, R. H. (1975). The methylation state of poly A-containing messenger RNA from cultured hamster cells. Nucleic Acids Res, 2(10), 1653–1668. 10.1093/nar/2.10.1653

Frost, B., Bardai, F. H., & Feany, M. B. (2016). Lamin Dysfunction Mediates Neurodegeneration in Tauopathies. Curr Biol, 26(1), 129–136. 10.1016/j.cub.2015.11.039

Frost, B., Hemberg, M., Lewis, J., & Feany, M. B. (2014). Tau promotes neurodegeneration through global chromatin relaxation. Nat Neurosci, 17(3), 357–366. 10.1038/nn.3639

Gokool, A., Loy, C. T., Halliday, G. M., & Voineagu, I. (2020). Circular RNAs: The Brain Transcriptome Comes Full Circle. Trends Neurosci, 43(10), 752–766. 10.1016/j.tins.2020.07.007

Gruner, H., Cortés-López, M., Cooper, D. A., Bauer, M., & Miura, P. (2016). CircRNA accumulation in the aging mouse brain. Sci Rep, 6, 38907. 10.1038/srep38907

Han, M., Liu, Z., Xu, Y., Liu, X., Wang, D., Li, F., … Bi, J. (2020). Abnormality of m6A mRNA Methylation Is Involved in Alzheimer’s Disease. Front Neurosci, 14, 98. 10.3389/fnins.2020.00098

Hanan, M., Simchovitz, A., Yayon, N., Vaknine, S., Cohen-Fultheim, R., Karmon, M., … Kadener, S. (2020). A Parkinson’s disease CircRNAs Resource reveals a link between circSLC8A1 and oxidative stress. EMBO Mol Med, 12(11), e13551. 10.15252/emmm.202013551

Hansen, T. B., Jensen, T. I., Clausen, B. H., Bramsen, J. B., Finsen, B., Damgaard, C. K., & Kjems, J. (2013). Natural RNA circles function as efficient microRNA sponges. Nature, 495(7441), 384–388. 10.1038/nature11993

Houseley, J. M., Garcia-Casado, Z., Pascual, M., Paricio, N., O’Dell, K. M., Monckton, D. G., & Artero, R. D. (2006). Noncanonical RNAs from transcripts of the Drosophila muscleblind gene. J Hered, 97(3), 253–260. 10.1093/jhered/esj037

Hsieh, Y. C., Guo, C., Yalamanchili, H. K., Abreha, M., Al-Ouran, R., Li, Y., … Shulman, J. M. (2019). Tau-Mediated Disruption of the Spliceosome Triggers Cryptic RNA Splicing and Neurodegeneration in Alzheimer’s Disease. Cell Rep, 29(2), 301–316 e310. 10.1016/j.celrep.2019.08.104

Huang, H., Camats-Perna, J., Medeiros, R., Anggono, V., & Widagdo, J. (2020). Altered Expression of the m6A Methyltransferase METTL3 in Alzheimer’s Disease. eNeuro, 7(5). 10.1523/eneuro.0125-20.2020

Hutton, M., Lendon, C. L., Rizzu, P., Baker, M., Froelich, S., Houlden, H., … Heutink, P. (1998). Association of missense and 5’-splice-site mutations in tau with the inherited dementia FTDP-17. Nature, 393(6686), 702–705. 10.1038/31508

Jankowsky, J. L., & Zheng, H. (2017). Practical considerations for choosing a mouse model of Alzheimer’s disease. Mol Neurodegener, 12(1), 89. 10.1186/s13024-017-0231-7

Janssen, J. C., Warrington, E. K., Morris, H. R., Lantos, P., Brown, J., Revesz, T., … Rossor, M. N. (2002). Clinical features of frontotemporal dementia due to the intronic tau 10(+16) mutation. Neurology, 58(8), 1161–1168. 10.1212/wnl.58.8.1161

Jiang, L., Lin, W., Zhang, C., Ash, P. E. A., Verma, M., Kwan, J., … Wolozin, B. (2021). Interaction of tau with HNRNPA2B1 and N 6-methyladenosine RNA mediates the progression of tauopathy. Mol Cell, 81(20), 4209–4227.e4212. 10.1016/j.molcel.2021.07.038

Jiang, X., Liu, B., Nie, Z., Duan, L., Xiong, Q., Jin, Z., … Chen, Y. (2021). The role of m6A modification in the biological functions and diseases. Signal Transduct Target Ther, 6(1), 74. 10.1038/s41392-020-00450-x

Kanadia, R. N., Johnstone, K. A., Mankodi, A., Lungu, C., Thornton, C. A., Esson, D., … Swanson, M. S. (2003). A muscleblind knockout model for myotonic dystrophy. Science, 302(5652), 1978–1980. 10.1126/science.1088583

Karch, C. M., Kao, A. W., Karydas, A., Onanuga, K., Martinez, R., Argouarch, A., … Group, T. C. S. C. (2019). A Comprehensive Resource for Induced Pluripotent Stem Cells from Patients with Primary Tauopathies. Stem Cell Reports, 13(5), 939–955. 10.1016/j.stemcr.2019.09.006

Khurana, V. (2008). Modeling Tauopathy in the fruit fly Drosophila melanogaster. J Alzheimers Dis, 15(4), 541–553. 10.3233/jad-2008-15403

Khurana, V., Merlo, P., DuBoff, B., Fulga, T. A., Sharp, K. A., Campbell, S. D., … Feany, M. B. (2012). A neuroprotective role for the DNA damage checkpoint in tauopathy. Aging Cell, 11(2), 360–362. 10.1111/j.1474-9726.2011.00778.x

Knopman, D. S., Amieva, H., Petersen, R. C., Chételat, G., Holtzman, D. M., Hyman, B. T., … Jones, D. T. (2021). Alzheimer disease. Nat Rev Dis Primers, 7(1), 33. 10.1038/s41572-021-00269-y

Knupp, D., & Miura, P. (2018). CircRNA accumulation: A new hallmark of aging? Mech Ageing Dev, 173, 71–79. 10.1016/j.mad.2018.05.001

Koren, S. A., Galvis-Escobar, S., & Abisambra, J. F. (2020). Tau-mediated dysregulation of RNA: Evidence for a common molecular mechanism of toxicity in frontotemporal dementia and other tauopathies. Neurobiol Dis, 141, 104939. 10.1016/j.nbd.2020.104939

Kyrylkova, K., Kyryachenko, S., Leid, M., & Kioussi, C. (2012). Detection of apoptosis by TUNEL assay. Methods Mol Biol, 887, 41–47. 10.1007/978-1-61779-860-3_5

Lence, T., Soller, M., & Roignant, J. Y. (2017). A fly view on the roles and mechanisms of the m. RNA Biol, 14(9), 1232–1240. 10.1080/15476286.2017.1307484

Li, L. B., Yu, Z., Teng, X., & Bonini, N. M. (2008). RNA toxicity is a component of ataxin-3 degeneration in Drosophila. Nature, 453(7198), 1107–1111. 10.1038/nature06909

Lin, J. J. (1981). Monoclonal antibodies against myofibrillar components of rat skeletal muscle decorate the intermediate filaments of cultured cells. Proc Natl Acad Sci U S A, 78(4), 2335–2339. 10.1073/pnas.78.4.2335

Mahoney, R., Ochoa Thomas, E., Ramirez, P., Miller, H. E., Beckmann, A., Zuniga, G., … Frost, B. (2020). Pathogenic Tau Causes a Toxic Depletion of Nuclear Calcium. Cell Rep, 32(2), 107900. 10.1016/j.celrep.2020.107900

Memczak, S., Jens, M., Elefsinioti, A., Torti, F., Krueger, J., Rybak, A., … Rajewsky, N. (2013). Circular RNAs are a large class of animal RNAs with regulatory potency. Nature, 495(7441), 333–338. 10.1038/nature11928

Memczak, S., Papavasileiou, P., Peters, O., & Rajewsky, N. (2015). Identification and Characterization of Circular RNAs As a New Class of Putative Biomarkers in Human Blood. PLoS One, 10(10), e0141214. 10.1371/journal.pone.0141214

Meng, S., Zhou, H., Feng, Z., Xu, Z., Tang, Y., Li, P., & Wu, M. (2017). CircRNA: functions and properties of a novel potential biomarker for cancer. Mol Cancer, 16(1), 94. 10.1186/s12943-017-0663-2

Miller, J. W., Urbinati, C. R., Teng-Umnuay, P., Stenberg, M. G., Byrne, B. J., Thornton, C. A., & Swanson, M. S. (2000). Recruitment of human muscleblind proteins to (CUG)(n) expansions associated with myotonic dystrophy. EMBO J, 19(17), 4439–4448. 10.1093/emboj/19.17.4439

Montalbano, M., McAllen, S., Puangmalai, N., Sengupta, U., Bhatt, N., Johnson, O. D., … Kayed, R. (2020). RNA-binding proteins Musashi and tau soluble aggregates initiate nuclear dysfunction. Nat Commun, 11(1), 4305. 10.1038/s41467-020-18022-6

Osouda, S., Nakamura, Y., de Saint Phalle, B., McConnell, M., Horigome, T., Sugiyama, S., … Furukawa, K. (2005). Null mutants of Drosophila B-type lamin Dm(0) show aberrant tissue differentiation rather than obvious nuclear shape distortion or specific defects during cell proliferation. Dev Biol, 284(1), 219–232. 10.1016/j.ydbio.2005.05.022

Pamudurti, N. R., Bartok, O., Jens, M., Ashwal-Fluss, R., Stottmeister, C., Ruhe, L., … Kadener, S. (2017). Translation of CircRNAs. Mol Cell, 66(1), 9–21.e27. 10.1016/j.molcel.2017.02.021

Pamudurti, N. R., Patop, I. L., Krishnamoorthy, A., Bartok, O., Maya, R., Lerner, N., … Kadener, S. (2022). circMbl functions in cis and in trans to regulate gene expression and physiology in a tissue-specific fashion. Cell Rep, 39(4), 110740. 10.1016/j.celrep.2022.110740

Paonessa, F., Evans, L. D., Solanki, R., Larrieu, D., Wray, S., Hardy, J., … Livesey, F. J. (2019). Microtubules Deform the Nuclear Membrane and Disrupt Nucleocytoplasmic Transport in Tau-Mediated Frontotemporal Dementia. Cell Rep, 26(3), 582–593 e585. 10.1016/j.celrep.2018.12.085

Paşca, A. M., Sloan, S. A., Clarke, L. E., Tian, Y., Makinson, C. D., Huber, N., … Paşca, S. P. (2015). Functional cortical neurons and astrocytes from human pluripotent stem cells in 3D culture. Nat Methods, 12(7), 671–678. 10.1038/nmeth.3415

Perry, R. P., Kelley, D. E., Friderici, K., & Rottman, F. (1975). The methylated constituents of L cell messenger RNA: evidence for an unusual cluster at the 5’ terminus. Cell, 4(4), 387–394. 10.1016/0092-8674(75)90159-2

Poorkaj, P., Bird, T. D., Wijsman, E., Nemens, E., Garruto, R. M., Anderson, L., … Schellenberg, G. D. (1998). Tau is a candidate gene for chromosome 17 frontotemporal dementia. Ann Neurol, 43(6), 815–825. 10.1002/ana.410430617

Rudnicki, D. D., Holmes, S. E., Lin, M. W., Thornton, C. A., Ross, C. A., & Margolis, R. L. (2007). Huntington’s disease--like 2 is associated with CUG repeat-containing RNA foci. Ann Neurol, 61(3), 272–282. 10.1002/ana.21081

Salzman, J., Chen, R. E., Olsen, M. N., Wang, P. L., & Brown, P. O. (2013). Cell-type specific features of circular RNA expression. PLoS Genet, 9(9), e1003777. 10.1371/journal.pgen.1003777

Sellier, C., Rau, F., Liu, Y., Tassone, F., Hukema, R. K., Gattoni, R., … Charlet-Berguerand, N. (2010). Sam68 sequestration and partial loss of function are associated with splicing alterations in FXTAS patients. EMBO J, 29(7), 1248–1261. 10.1038/emboj.2010.21

Shao, Y., & Chen, Y. (2016). Roles of Circular RNAs in Neurologic Disease. Front Mol Neurosci, 9, 25. 10.3389/fnmol.2016.00025

Shafik, A. M., Zhang, F., Guo, Z., Dai, Q., Pajdzik, K., Li, Y., Kang, Y., Yao, B., Wu, H., He, C., Allen, E. G., Duan, R., & Jin, P. (2021, Jan 05). N6-methyladenosine dynamics in neurodevelopment and aging, and its potential role in Alzheimer’s disease. Genome Biol, 22(1), 17. 10.1186/s13059-020-02249-z

Sohn, C., Ma, J., Ray, W. J., & Frost, B. (2023). Pathogenic tau decreases nuclear tension in cultured neurons. Front Aging, 4, 1058968. 10.3389/fragi.2023.1058968

Speese, S. D., Ashley, J., Jokhi, V., Nunnari, J., Barria, R., Li, Y., … Budnik, V. (2012). Nuclear envelope budding enables large ribonucleoprotein particle export during synaptic Wnt signaling. Cell, 149(4), 832–846. 10.1016/j.cell.2012.03.032

Spillantini, M. G., Murrell, J. R., Goedert, M., Farlow, M. R., Klug, A., & Ghetti, B. (1998). Mutation in the tau gene in familial multiple system tauopathy with presenile dementia. Proc Natl Acad Sci U S A, 95(13), 7737–7741. 10.1073/pnas.95.13.7737

Tanabe, L. M., Liang, C. C., & Dauer, W. T. (2016). Neuronal Nuclear Membrane Budding Occurs during a Developmental Window Modulated by Torsin Paralogs. Cell Rep, 16(12), 3322–3333. 10.1016/j.celrep.2016.08.044

Wang, Y., & Wang, Z. (2015). Efficient backsplicing produces translatable circular mRNAs. RNA, 21(2), 172–179. 10.1261/rna.048272.114

Wittmann, C. W., Wszolek, M. F., Shulman, J. M., Salvaterra, P. M., Lewis, J., Hutton, M., & Feany, M. B. (2001). Tauopathy in Drosophila: neurodegeneration without neurofibrillary tangles. Science, 293(5530), 711–714. 10.1126/science.1062382

Yang, Y., Fan, X., Mao, M., Song, X., Wu, P., Zhang, Y., … Wang, Z. (2017). Extensive translation of circular RNAs driven by N. Cell Res, 27(5), 626–641. 10.1038/cr.2017.31

Zhou, C., Molinie, B., Daneshvar, K., Pondick, J. V., Wang, J., Van Wittenberghe, N., … Mullen, A. C. (2017). Genome-Wide Maps of m6A circRNAs Identify Widespread and Cell-Type-Specific Methylation Patterns that Are Distinct from mRNAs. Cell Rep, 20(9), 2262–2276. 10.1016/j.celrep.2017.08.027

Zhou, M., Xiao, M. S., Li, Z., & Huang, C. (2021). New progresses of circular RNA biology: from nuclear export to degradation. RNA Biol, 18(10), 1365–1373. 10.1080/15476286.2020.1853977

Zuniga, G., Levy, S., Ramirez, P., De Mange, J., Gonzalez, E., Gamez, M., & Frost, B. (2022). Tau-induced deficits in nonsense-mediated mRNA decay contribute to neurodegeneration. Alzheimers Dement. 10.1002/alz.12653

